# Extracellular vesicle-mediated release of bis(monoacylglycerol)phosphate is regulated by LRRK2 and Glucocerebrosidase activity

**DOI:** 10.1101/2023.07.12.548710

**Authors:** Elsa Meneses-Salas, Moisés Castellá, Marianna Arnold, Frank Hsieh, Ruben Fernández-Santiago, Mario Ezquerra, Alicia Garrido, María-José Martí, Carlos Enrich, Suzanne R. Pfeffer, Kalpana Merchant, Albert Lu

## Abstract

The endolysosomal phospholipid, bis(monoacylglycerol)phosphate (BMP), is aberrantly elevated in the urine of Parkinson’s patients with mutations in genes encoding leucine-rich repeat kinase 2 (LRRK2) and glucocerebrosidase (GCase). Because BMP resides on and regulates the biogenesis of endolysosomal intralumenal membranes that become extracellular vesicles (EVs) upon release, we hypothesized that elevated urinary BMP may be driven by increased exocytosis of BMP-enriched EVs. To test this hypothesis, we analyzed BMP metabolism and EV-associated release of BMP in wild type (WT) and R1441G LRRK2-expressing mouse embryonic fibroblast (MEF) cells. Using immunofluorescence microscopy and transmission electron microscopy we detected structural alterations in endolysosomes and antibody-accessible BMP pool, indicating that mutant LRRK2 affects endolysosomal homeostasis. Biochemical analyses of isolated EV fractions confirmed the effect on endolysosomes by showing an increase in LAMP2-positive EVs in mutant cells, which was partially restored by LRRK2 kinase inhibition but further augmented, albeit to a more variable extent, by GCase inhibition. Using mass spectrometry, we detected an overall increase in total di-22:6-BMP and total di-18:1-BMP in cell lysates from mutant LRRK2 MEFs compared to WT cells. Inhibition of LRRK2 kinase partially restored cellular BMP levels, whereas inhibition of GCase further increased the BMP content. In isolated EVs from LRRK2 mutant cells, LRRK2 inhibition decreased BMP content whereas GCase inhibition tended to increase it. Using metabolic labeling experiments, we demonstrated that the increase in cellular BMP content is not due to an increase in BMP synthesis, even though we observed an increase in BMP synthesizing enzyme, CLN5, in LRRK2 mutant MEFs and patient-derived fibroblasts. Finally, pharmacological modulation of EV release and live total internal reflection fluorescence (TIRF) microscopy of endolysosomal exocytosis in human G2019S LRRK2 fibroblasts, further confirmed that BMP release is associated with EV secretion. Together, these results establish LRRK2 as a regulator of BMP levels in cells and its release through EVs and suggest that GCase activity further modulates this process in LRRK2 mutant cells. Mechanistic insights from these studies have implications for the potential use of BMP-positive EVs as a biomarker for Parkinson’s disease and associated treatments.

## INTRODUCTION

Variants in the gene encoding Leucine rich repeat kinase 2 (*LRRK2*) and β-glucocerebrosidase (GCase; *GBA1*) together constitute the largest contributors to familial and sporadic Parkinson’s disease (PD)^1^. Although G2019S is the most prevalent PD-associated *LRRK2* mutation, R1441G is more penetrant and is associated with an earlier disease onset^2–4^. A shared mechanism of pathogenesis among all PD-causing LRRK2 mutations is increased LRRK2 kinase activity, which leads to phosphorylation of its cognate Rab GTPase substrates, including Rab8, Rab10 and Rab12, that are key regulators of intracellular vesicle trafficking^5^. A prevailing hypothesis of pathogenesis in LRRK2 PD cases is that LRRK2 phosphorylation of Rabs results in defective autophagy and endolysosomal homeostasis and defective ciliogenesis in the nigrostriatal circuit^5,6^.

*GBA1* encodes a lysosomal hydrolase, glucocerebrosidase (GCase), essential for glycosphingolipid degradation. Homozygous loss-of-function mutations in *GBA1* cause Gaucher’s disease, a lysosomal storage disorder characterized by cytotoxic accumulation of lipid substrates of GCase, glucosylceramide and glucosylsphingosine; heterozygous carriers have a 5-10 fold increased risk of PD ^7,8^. The predominant hypothesis for the pathogenic mechanism of *GBA1* variants is related to aberrant aggregation and clearance of alpha-synuclein in a feed-forward cascade^9,10^; a hypothesis being tested clinically using GCase activators. Additionally, we and others previously showed that GCase enzymatic function is modulated by LRRK2 kinase activity^11,12^. Altogether, it is possible that dysregulation of the endolysosomal biology is associated with pathogenic variants of both *LRRK2* and *GBA1*^6,13^.

Bis(monoacylglycerol)phosphate (BMP, also known as LBPA or Lysobisphosphatidic acid) is an atypical, negatively-charged glycerophospholipid important for maintaining endolysosomal homeostasis^14^. BMP resides primarily in endolysosomal intralumenal vesicles (ILVs) where it participates in multiple processes, including cholesterol egress, exosome biogenesis and lipid catabolism^14,15^. Of note, BMP acts together with Saposin-C as enzymatic cofactors of GCase^16,17^. Multiple BMP isomers are present in cells, based on differences in their fatty acyl chains and their positions relative to the sn-2 and sn-3 carbon atoms on each of its two glycerol backbones^14,15^; the sn-2:sn-2’ isomer is the proposed active form^18,19^. Dioleoyl-BMP (di-18:1-BMP) is the predominant form in numerous cell lines analyzed^20–23^ and didocosahexaenoyl-BMP (di-22:6-BMP) is one of the most abundant species in the brain^24,25^. Fatty acyl composition is thought to influence BMP biophysical properties and function^26^.

We and others recently reported elevated levels of total di-18:1-BMP and total di-22:6-BMP species in the urine of carriers of PD-associated *LRRK2* and *GBA1* mutations^27–29^. Although we did not observe a correlation between urinary BMP and PD progression^28^, these studies underscore BMP’s utility as a patient enrichment and target modulation biomarker in therapeutic trials^30^. Based on our previous findings, in the present study we focused on these two BMP species.

Given the specific intracellular localization of BMP (in ILVs; exosomes when released) and its proposed roles in exosome biogenesis^19^, we hypothesized that its increased levels in PD biofluids may be a consequence of an increase in the secretion of BMP containing extracellular vesicle (EV). To gain mechanistic insight into the regulation and biological significance of BMP release, we monitored BMP release in EVs and performed kinetic analyses of BMP biosynthesis in mouse embryonic fibroblasts (MEFs) from wild-type (WT) or R1441G LRRK2 knock-in mice. Moreover, endolysosomal exocytosis was visualized and quantified in skin fibroblasts derived from both healthy donors and G2019S LRRK2 PD-patients. In addition, we examined the regulation of BMP by a LRRK2 kinase inhibitor, MLi-2, and a GCase inhibitor, conduritol β-epoxide (CBE), to gain insights into the role of BMP in potential therapeutic vs. pathophysiological responses, respectively. Our data indicate that LRRK2 kinase activity modulate BMP release in EVs and further suggest that GCase function may contribute to this process.

## RESULTS

### Analysis of BMP-positive endolysosomes in R1441G LRRK2 mouse embryonic fibroblasts

We first performed immunofluorescence microscopy of BMP-positive endolysosomes in both WT and R1441G LRRK2 knock-in MEFs. The overall antibody-accessible BMP pool was significantly decreased in mutant LRRK2-expressing cells compared to WT (Figure 1A, B). In contrast, endolysosomal LAMP2 fluorescence was higher in mutant LRRK2 MEFs (Figure 1A, C). Previous studies found structural defects in endolysosomes of different cell types harboring pathogenic LRRK2 mutations^31–33^. Since BMP is specifically enriched in intra-luminal vesicles (ILVs) in the endolysosomes where it regulates important catabolic reactions, we examined multivesicular endosome (MVE) morphology of these cells. In agreement with previous findings^33^, transmission electron microscopy revealed decreased MVE area in R1441G LRRK2 MEFs (average area= 0.11 µm^2^) compared to WT MEFs (average area= 0.27 µm^2^) (Figure 1D, E). In addition, the overall endolysosomal ILV number was diminished in the R1441G LRRK2 cell line (Figure 1F). These data suggest that the pathogenic LRRK2 mutation alters BMP-positive endolysosome morphology and ILV content.

**Figure 1.**
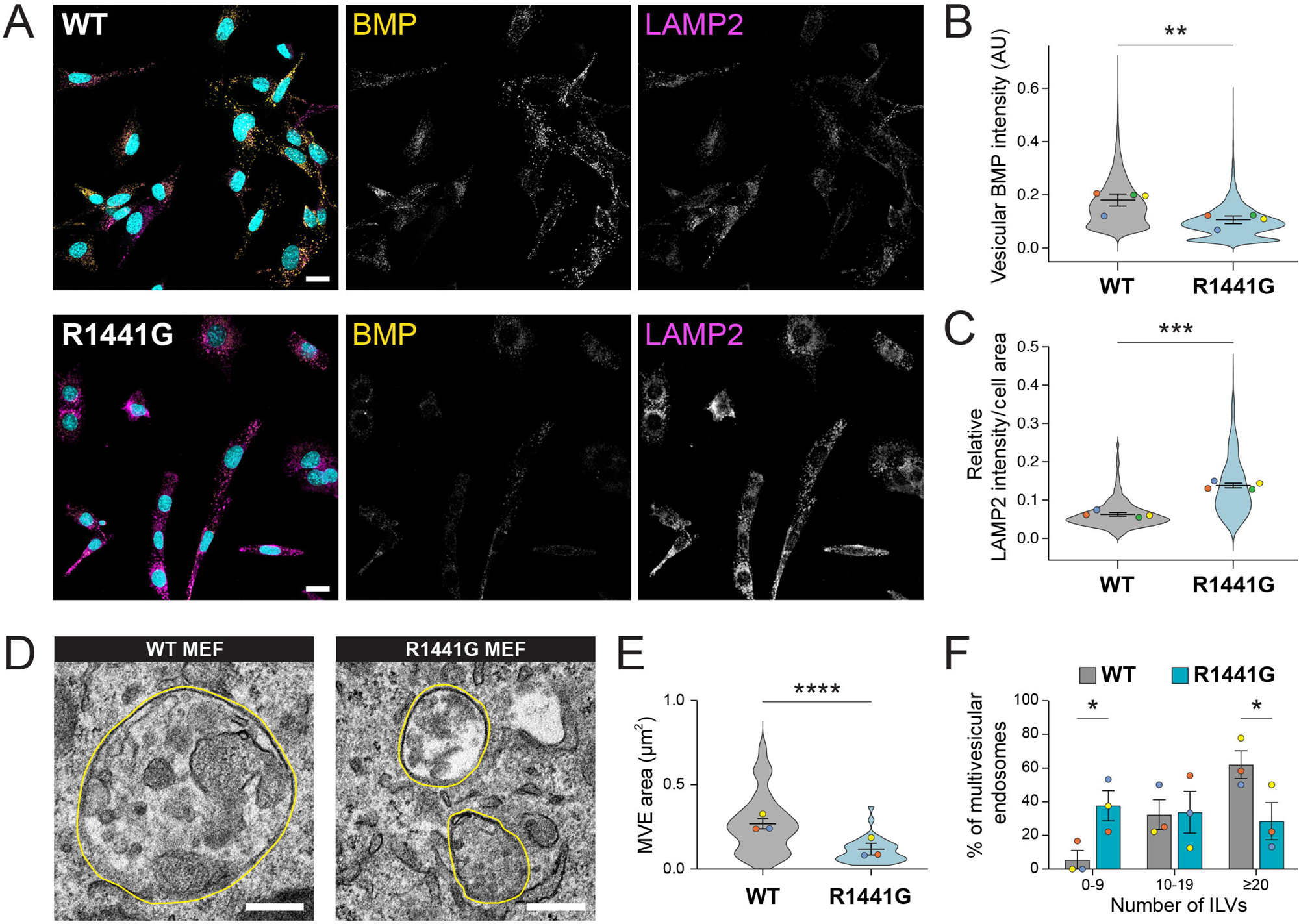
Alterations in antibody-accessible BMP and endolysosomal morphology in R1441G LRRK2 MEF cells. **(A)** Confocal microscopy of endogenous BMP (green) and LAMP2 (red) immunofluorescence in WT and R1441G LRRK2 MEFs. Scale bar: 20 µm. **(B-C)** Quantification of vesicular BMP intensity (**B**) and LAMP2 relative intensity (**C**) per cell area. Colored dots represent mean value from 4 independent experiments and violin plots show the distribution of individual cell data. Significance determined by two-tailed paired *t* test **p<0.01, ***p<0.001. **(D)** Representative transmission electron microscopy (TEM) images of Multivesicular endosomes (MVE) from WT and R1441G LRRK2 MEFs. MVB periphery highlighted in yellow. Scale bar: 250 nm. **(E)** MVE area (µm^2^) quantification in WT and R1441G LRRK2 mutant cells. Colored dots represent mean values from 3 independent experiments and violin plots show the distribution of individual cell data (35-45 cells/group). **(F)** Quantification of Intraluminal Vesicles (ILVs) per MVE in WT and R1441G LRRK2 MEF cells. The number of ILVs per MVE are binned in three groups and plotted as a percentage of MVE from the total population of each experiment independently. Data from 3 independent experiments (mean ± SEM). Significance determined by two-tailed unpaired *t* test **(E)** and ordinary two-way ANOVA, uncorrected Fisher’s LSD **(F)** *p<0.05, ****p<0.0001.

### Investigating the impact of LRRK2 and GCase activities on extracellular vesicle release modulation

Given the previously described roles of BMP in ILV/exosome biogenesis, we investigated whether the alterations observed in antibody-accessible BMP and MVE ILV number in LRRK2 mutant MEFs could be explained by changes in EV release. We isolated and characterized EVs from both WT and R1441G MEFs and also assessed the effects of LRRK2 and GCase pharmacological inhibitors, MLi-2 and CBE, respectively on EV content and number. Inhibition of LRRK2 kinase and GCase enzymatic activities was confirmed, respectively, by monitoring phospho-Rab10 levels in whole cell lysates and performing a fluorescence-based GCase activity assay in cells (Figure 2A, B). Consistent with our immunofluorescence data, upregulation of LAMP2 was observed in R1441G LRRK2 whole cell lysates (WCL; Figure 2 A, C, D and E), as previously reported in LRRK2 G2019S knock-in mouse brain^34^. MLi-2 treatment for 48 hours partially reversed this phenotype (Figure 2 A, C, E), suggesting that aberrant LRRK2 kinase activity influences endolysosomal homeostasis.

**Figure 2.**
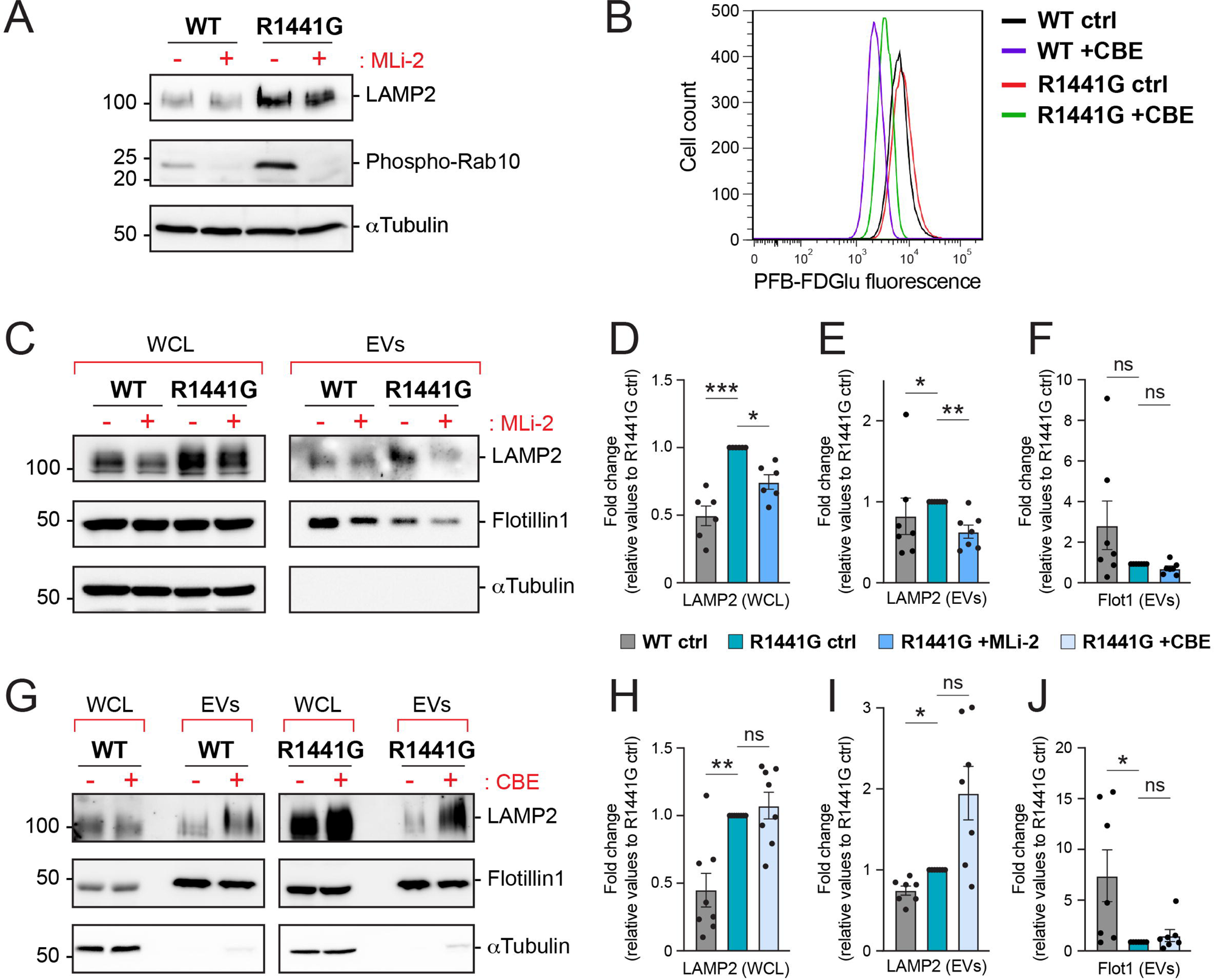
LRRK2 and GCase activities modulate extracellular vesicle production. **(A)** Whole cell lysates from WT and R1441G LRRK2 MEF cells treated with 200nM MLi-2 for 24h were analyzed by immunoblotting. Representative images of LAMP2, phospho-Rab10 and α-Tubulin levels are shown. Molecular weight marker mobility is shown in kDa. **(B)** Flow cytometry measurement of GCase activity using PFB-FDGlu fluorescent GCase substrate in WT and R1441G LRRK2 mutant MEF cells treated with 300µM CBE for 24h. **(C and G)** Whole cell lysates (WCL) and isolated extracellular vesicles (EVs) from WT and R1441G LRRK2 mutant MEF cells treated with 200nM MLi-2 **(C)** or 300µM CBE **(G)** for 48h were analyzed by immunoblotting. Representative images of LAMP2, Flotillin-1 and α-Tubulin levels are shown. Molecular weight marker mobility is shown in kDa. Immunoblots for LAMP2 and Flotillin-1 in EV fractions required longer exposure times to visualize clear signals across all conditions. **(D-F and H-J)** Quantification of LAMP2 and Flotillin-1 levels relative to R1441G LRRK2 MEF cells in whole cell lysates (WCL). **(D and H)** and isolated EVs **(E,F,I and J).** Data from 6-8 independent experiments (mean ± SEM). Significance determined by Kruskal-Wallis test followed by an uncorrected Dunn’s post hoc test compared to R1441G LRRK2 control *p<0.05, **p<0.01, ****p<0.0001; ns, not significant.

We next analyzed the profiles of isolated EV fractions from both wild-type (WT) and R1441G LRRK2 mutant MEFs. Both LAMP2 and Flotillin-1 were assessed to explore alterations in release of EV subpopulations; LAMP2 is enriched in ILV-derived EVs while Flotillin-1 is also seen in plasma membrane-derived ectosomes that reflect outward budding of the plasma membrane^35–37^. Biochemical analysis revealed an elevation of LAMP2, but not Flotillin-1, in EVs derived from R1441G LRRK2 MEFs compared to those from WT MEFs (Figure 2C, F, G). Upon MLi-2 treatment, a modest but significant decrease in both EV-associated protein markers was observed in isolated EV fractions from mutant LRRK2 MEFs (Figure 2C, F, G). In contrast, inhibition of GCase activity in mutant LRRK2 cells led to an increase in LAMP2, but not Flotillin-1, in isolated EVs compared to those derived from WT MEFs (Figure 2C, F, G). Finally, analysis of WT cell-derived EVs revealed no major differences in EV marker levels between untreated (control) and MLi-2- or CBE-treated conditions (Supplemental Figure 1A-C).

To complement these data, we conducted nanoparticle tracking analysis (NTA). Isolated EV fractions from WT and R1441G LRRK2 cells exhibited comparable particle size distributions (Supplemental Figure 1D). Treatments with MLi-2 and CBE yielded measurable quantitative changes in EV concentrations that, despite not reaching statistical significance, gave similar trends as those seen in our biochemical analyses (Supplemental Figure 1E), potentially reflecting the inherent variability of NTA due to its inability to distinguish EVs from non-vesicular particles^38,39^. For the R1441G MEF cells, MLi-2 decreased EV concentration while CBE increased EV particles per ml, in agreement with the effects observed in our biochemical analysis.

Altogether, these results suggest that EV secretion is influenced by LRRK2 kinase and, to a more variable degree, by GCase hydrolase activities; while MLi-2 treatment decreases EV release, CBE appears to increase it.

### Targeted BMP lipid analysis in intact cells and extracellular vesicles

Ultra-performance liquid chromatography-tandem mass spectrometry (UPLC-MS/MS) was used to measure quantitatively the impact of LRRK2 and GCase activities on BMP isoform abundance in cells and isolated EV fractions. Remarkably, an overall increase in total BMP isoforms was detected in mutant LRRK2 MEF cell lysates (Figure 3A-D), with di-22:6-BMP and di-18:1-BMP as the major species. It is important to note that mass spectrometry-based methods detect total BMP content while antibody staining (Figure 1A, B) only detects so-called “antibody-accessible” BMP^40^. The mass spectrometry-based increase in total BMP in LRRK2 mutant cells indicates that this pool is less antibody-accessible than that present in wild type cells. Alternately, the anti-BMP antibody may be less specific and detect other analytes.

**Figure 3.**
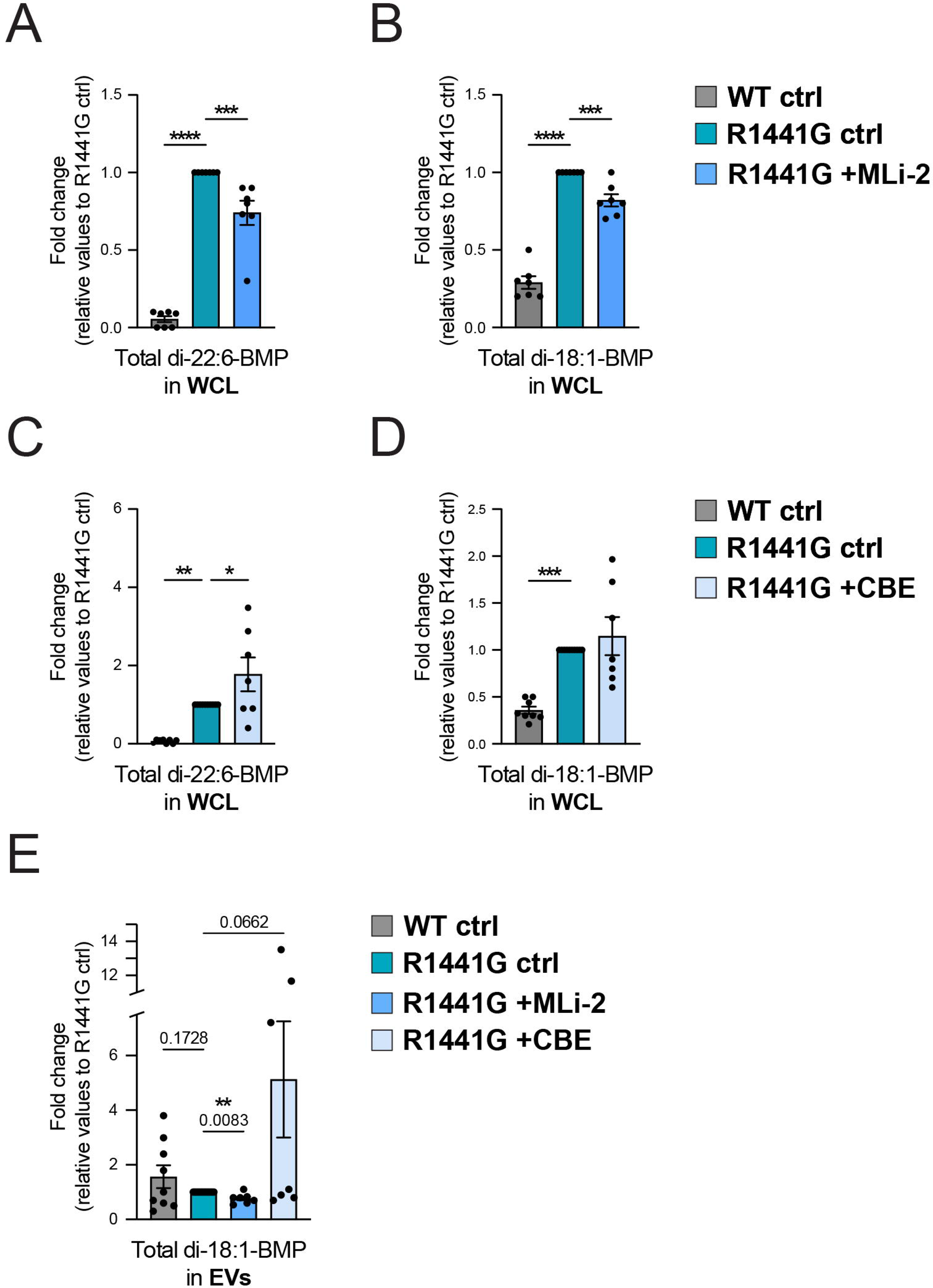
Targeted lipid pathway analysis of BMP abundance in cellular and isolated EV fractions. **(A-D)** UPLC-MS/MS determination of BMP isoforms normalized to protein content from cells treated with 200nM MLi-2 **(A-B)** or 300µM CBE **(C-D)** for 48h. Data shown as fold change relative to untreated R1441G LRRK2 MEF cells. Only BMP isoforms that were detected are shown. **(E)** UPLC-MS/MS determination of BMP isoforms normalized to protein content in EVs isolated from cells treated with 200nM MLi-2 or 300µM CBE for 48h. Only BMP isoforms that were detected are shown. Data from 3-6 independent experiments (mean ± SEM). Significance determined by ordinary one-way ANOVA, uncorrected Fisher’s LSD **(A-D)** and one-way ANOVA with the Geisser-Greenhouse correction, uncorrected Fisher’s LSD **(E)** *p<0.05, **p<0.01, ***p<0.001, ****p<0.0001.

In R1441G cells, LRRK2 inhibition for 48 hours with MLi-2 decreased total di-22:6-BMP and total di-18:1-BMP levels by ∼20% (Figure 3A and B). Conversely, inhibition of GCase increased total BMP levels, reaching significance for di-22:6-BMP levels compared with untreated, R1441G LRRK2 cells (Figure 3C). No statistically significant differences in intracellular BMP levels were observed in WT LRRK2 MEFs upon LRRK2 or GCase inhibition (Supplemental Figure 1F, G), suggesting a dominant role of mutant LRRK2 activity in BMP regulation. Overall, di-oleoyl (18:1)- and di-docosahexaenoyl (22:6)-BMP species were the most abundant in MEFs, whereas other isoforms, such as di-arachidonoyl (20:4)- and dilinoleoyl (18:2)-BMP, were present at lower levels but were also consistently elevated in mutant LRRK2 MEFs (Supplemental Figure 1H). These data suggest that there are LRRK2-independent clonal differences leading to differences in basal BMP content between WT and mutant MEF cells; however, such clonal variation does not impact the effect of MLi-2 or CBE treatment in R1441G cells.

Analysis of isolated EV fractions detected only the di-18:1-BMP isoform, with no evidence of di-22:6-BMP. Although a trend toward higher di-18:1-BMP levels was observed in EVs derived from WT cells compared to those from mutant LRRK2 cells, this difference was not statistically significant (Figure 3E). Treatment of LRRK2 R1441G MEFs with MLi-2 resulted in a significant partial decrease in EV-associated total BMP (Figure 3E), consistent with previous observations in non-human primates and PD mouse models that showed decreased extracellular urinary BMP upon LRRK2 kinase pharmacological inhibition^41–43^. In contrast, inhibition of GCase activity yielded an opposite, albeit not significant, trend (Figure 3E). These latter findings may be explained by the observation that GCase inhibition by CBE was less pronounced in LRRK2 R1441G cells compared to WT cells under identical concentration and treatment duration conditions (Figure 2B). Finally, no significant differences in EV-associated BMP abundance between untreated (control) and MLi-2- or CBE-treated WT LRRK2 MEFs were observed (Supplemental Figure 1I).

In addition to analyzing BMP, we also examined GCase lipid substrates in both cells and isolated EVs. Targeted quantification of sphingolipid species revealed elevated GCase substrate abundance in both CBE-treated WT and mutant LRRK2 cells, validating the treatment paradigm used in this study. Interestingly, CBE-mediated accumulation of GCase substrates was more evident in R1441G LRRK2 than in WT cells (Supplemental Figure 2A). This suggests that, despite a less pronounced direct effect on GCase activity (Figure 2B), the R1441G mutation might contribute to broader endolysosome dysfunction, which could alter the dynamics of substrate accumulation. On the other hand, analysis of isolated EV fractions revealed lower levels of glucosylceramide, galactosylceramide, and glucosylsphingosine in EVs from R1441G LRRK2 cells compared to those from WT MEFs, but no significant differences were detected between control or CBE-treated conditions independent of LRRK2 mutation status (except for glucosylsphingosine, which showed an increase in WT-EVs upon GCase inhibition; Supplemental Figure 2B).

Altogether, given that BMP is specifically enriched in ILVs (which become exosomes upon release), the data presented above support our biochemical analysis (Figure 2C, E, G, I) and suggest that LRRK2 activity regulates BMP release in association with LAMP2-positive exosomes, whereas GCase activity appears to have a more variable effect under the tested conditions.

### BMP biosynthesis is not influenced by LRRK2 kinase and GCase activities

To rule out the possibility that differences in cellular and EV-associated BMP levels following MLi-2 or CBE treatment of R1441G cells were caused by alterations in BMP metabolism rather than changes in membrane trafficking and EV release, we performed metabolic labeling experiments using heavy (H) isotope-labeled 22:6 and 18:1 fatty acids as BMP precursors.

Cells were pulsed for 25_min with heavy isotope BMP precursors and then washed and chased for different time points (Figure 4A). UPLC-MS/MS analysis allowed us to differentiate unlabeled versus semi-labeled (H,L’; only one heavy isotope-labeled fatty acid chain) or fully-labeled (H,H’; both fatty acid chains heavy isotope-labeled) BMP species (Figure 4A). Initial experiments were performed over 48h using mutant LRRK2 cells under control and LRRK2- or GCase-inhibition conditions; untreated WT cells were included for comparison. Strikingly, even at time 0 and throughout the length of the experiment, R1441G LRRK2 cells displayed substantially higher H,L’- and H,H’-22:6-BMP species than WT cells (left and middle plots in Figure 4 B). H,L’, but not H,H’ 18:1-BMP was detected, which appeared to increase to a greater extent in R1441G LRRK2 cells at least during the initial 8h of the experiment (right plot, Figure 4 B). Despite these differences between WT and mutant LRRK2 cells, neither inhibition of LRRK2 or GCase had a major impact in the kinetics of BMP synthesis or catabolism (Figure 4B).

**Figure 4.**
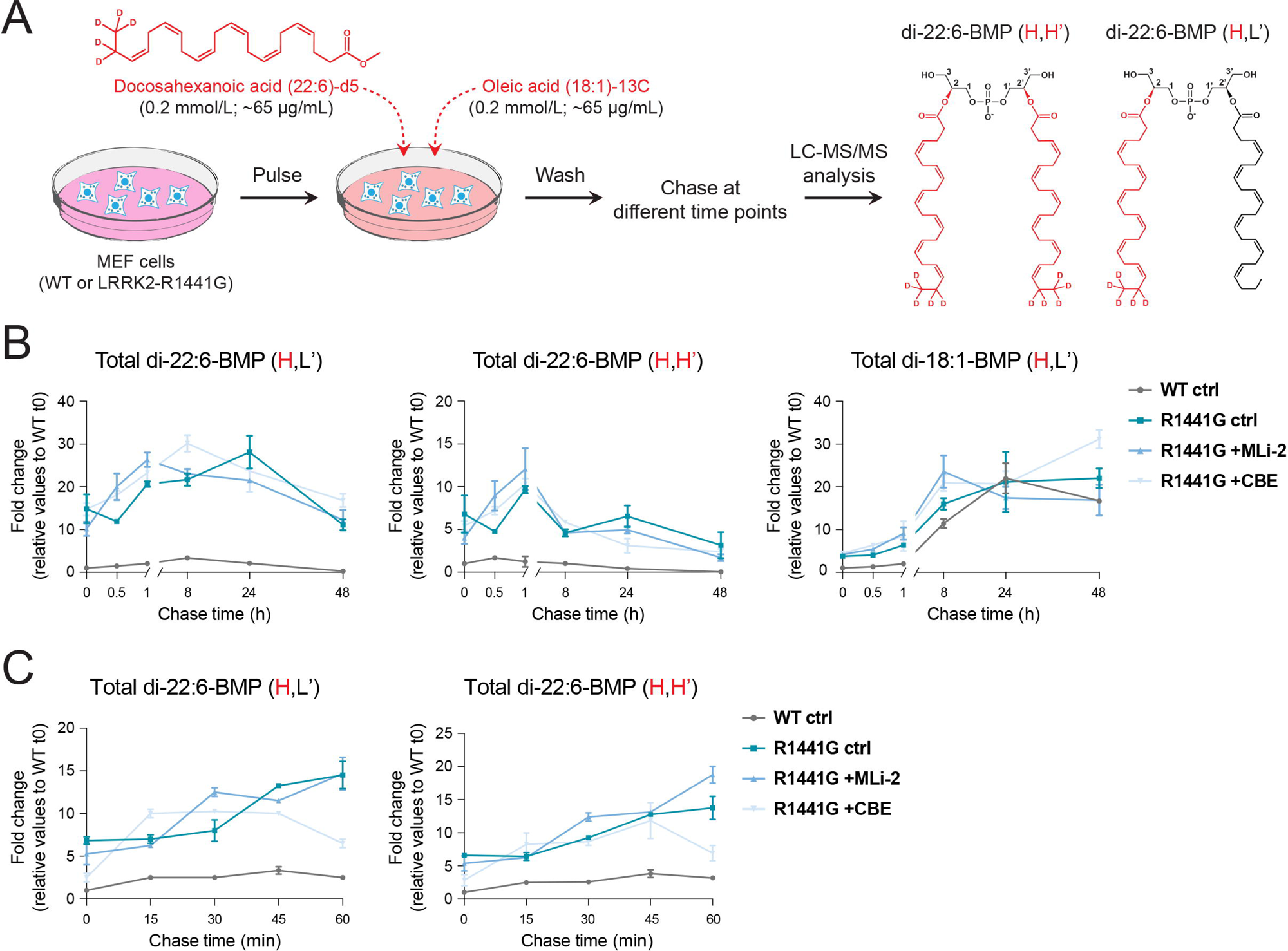
Inhibition of LRRK2 or GCase activities does not significantly impact BMP biosynthetic and catabolic rates. **(A)** Schematic representation of the BMP metabolic labeling protocol with deuterated docosahexaenoic acid and ^13^C-labeled oleic acid. WT and R1441G mutant MEF cells were incubated with a pulse of DHA-d5 / OA-^13^C for 20min, followed by a chase for different times. Cells were then collected for subsequent BMP lipidomic analysis. Structures of unlabeled (L) and isotope-labeled (H) fatty acids are shown in black or red, respectively. **(B-D)** UPLC-MS/MS determination of BMP isoforms normalized to protein content from WT MEF cells and R1441G LRRK2 mutant MEF cells ± MLi-2 (200nM) or CBE (300µM). Long **(B-C)** and short **(D)** chase time points shown as fold change relative to WT control MEF cells time 0. Only BMP isoforms that were detected are shown. Data from 3 replicated experiments (mean ± SEM).

In these experiments, semi-labeled BMP species gradually increased for up to 24 h. This likely reflects re-utilization of isotope-labeled lysophosphatidylglycerol (LPG) and fatty acid precursors generated after degradation of either H,L’ or H,H’ BMP the by endolysosomal hydrolase PLA2G15^44^. To better resolve initial BMP synthesis rates, we reduced the pulse labeling time and chase time to 60 min. In this set of experiments, only isotope-labeled 22:6-BMP species were detected (Figure 4C), possibly reflecting a preference for 22:6 BMP synthesis by MEFs. As before, even at time 0 and throughout the 60 min duration of the assay, 22:6-BMP levels were consistently higher in R1441G LRRK2 cells compared with WT MEFs. Again, no overall rate differences were seen between untreated and MLi-2- or CBE-treated cells. These experiments support the hypothesis that rather than altering BMP metabolic rates, LRRK2 and GCase activities influence EV-mediated BMP release.

### Increased BMP synthase protein expression in mutant LRRK2 cells

To further investigate the biological significance of BMP upregulation observed in mutant LRRK2 MEF cells (Figure 3A-D; Figure 4B, C), we examined the expression of CLN5, a key lysosomal enzyme in the BMP biosynthetic pathway^45,46^. Interestingly, biochemical analysis of total cell lysates revealed a significant fold-change increase in CLN5 protein levels in R1441G LRRK2 MEFs relative to WT cells (Figure 5A). Notably, 16 hour treatment with MLi-2 reduced CLN5 levels to a comparable fold-change extent (Figure 5B). To validate these findings in a human and disease-relevant context, we examined CLN5 protein expression in cultured fibroblasts derived from Parkinson’s disease (PD) patients carrying the G2019S LRRK2 mutation. Consistent with our observations in MEF cells, CLN5 protein levels were reduced following 16 h MLi-2 treatment in a concentration-dependent manner (Figure 5C). Notably, the maximal reduction was observed at the same 200nM MLi-2 concentration used in MEF cell experiments, a dose that also did not induce observable cytotoxic effects in human fibroblasts. Similar results were obtained in patient-derived fibroblasts harboring the R1441G LRRK2 mutation (Figure 5D). Altogether, these data suggest that LRRK2 kinase activity may regulate CLN5 protein expression.

**Figure 5.**
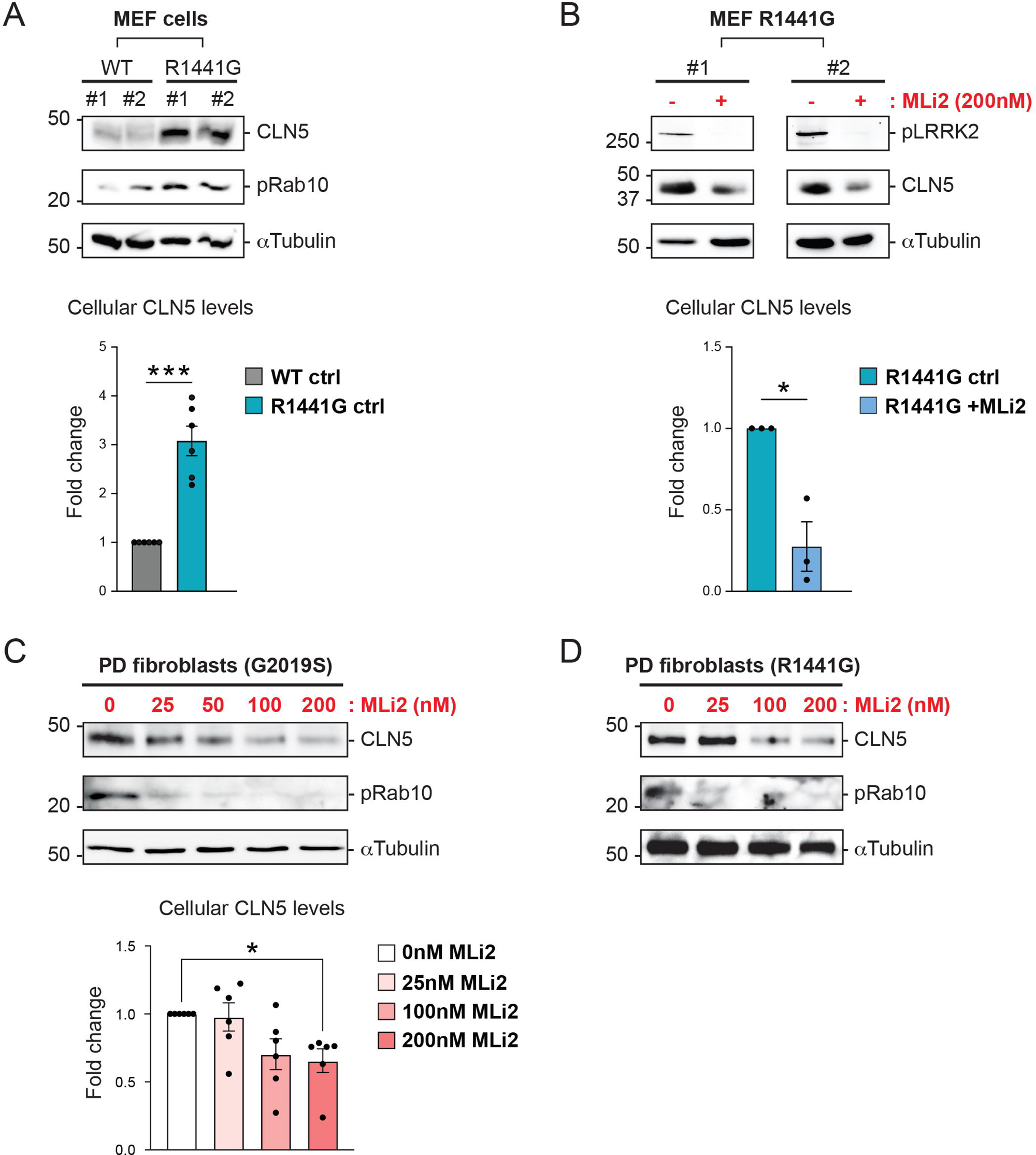
LRRK2 activity modulates CLN5 expression levels. **(A)** Whole cell lysates from WT and R1441G LRRK2 MEF cells were analyzed by immunoblotting. Representative immunoblots of CLN5, phospho-Rab10 (pRab10) and α-Tubulin are shown from two (#1 and #2) out of six independent experiments. Molecular weight marker mobility is shown in kDa. Plot at the bottom shows quantification of CLN5 immunoblot levels relative to WT and R1441G LRRK2 MEF cells**. (B)** Whole cell lysates from R1441G LRRK2 MEF cells treated with 200nM MLi-2 for 24h were analyzed by immunoblotting. Representative immunoblots of phosphor-LRRK2 (pLRRRK2), CLN5 and α-Tubulin levels are shown from two (#1 and #2) out of three independent experiments. Molecular weight marker mobility is shown in kDa. Plot at the bottom shows quantification of CLN5 immunoblot levels in whole cell lysates of R1441G LRRK2 MEF cells untreated or MLi-2-treated**. (C)** Whole cell lysates from G2019S LRRK2 patient-derived fibroblasts treated with indicated increasing MLi-2 concentrations for 24h were analyzed by immunoblotting. Immunoblots of CLN5, phospho-Rab10 (pRab10) and α-Tubulin levels are shown from one representative experiment (n=6 per condition, obtained from two independent replicate experiments using fibroblast cell lines derived from three different patients). Molecular weight marker mobility is shown in kDa. Plot at the bottom shows quantification of CLN5 immunoblot levels relative to G2019S LRRK2 patient-derived fibroblasts treated with MLi-2 at the indicated concentrations**. (D)** Whole cell lysates from R1441G LRRK2 patient-derived fibroblasts treated with indicated increasing MLi-2 concentrations for 24h were analyzed by immunoblotting. Immunoblots of CLN5, phospho-Rab10 (pRab10) and α-Tubulin levels are shown from one representative experiment. Significance in **(A)** and **(B)** determined by two-tailed, unpaired *t* test; significance in **(C)** determined by Dunnett’s One-way ANOVA test; *p<0.05, ***p<0.001.

The upregulation of CLN5 may be due to an overall upregulation of lysosomal enzymes as LAMP2 levels were also increased (Figure 2A, C, E). Moreover, the observed baseline differences in BMP, as documented in the flux study above (Figure 4B and C), could result from CLN5 upregulation. The lack of significant changes in the BMP synthesis rate (Figure 4B and C) suggests either a limitation in substrate availability or that CLN5 is operating at maximal capacity.

### BMP release is EV-mediated

Our data suggest that BMP is exocytosed in association with EVs and that LRRK2 and GCase activities modulate BMP secretion. To determine the magnitude of BMP release via EV secretion, we assessed the impact of pharmacological modulators of EV release. Treatment of WT MEFs with GW4869, a selective type 2-neutral sphingomyelinase inhibitor, decreased EV release as monitored by reduced levels of LAMP2 and Flotillin-1 in EV fractions (Figure 6A). Under these conditions, we observed a parallel decrease in exosomal BMP (Figure 6A, B). In contrast, enhancing EV release with bafilomycin A1 (B-A1), a pharmacological inhibitor of the endolysosomal proton pump V-ATPase that dramatically boosts EV exocytosis^36,47^, resulted in the opposite trend (Figure 6A, C): EV markers and BMP levels increased. Biochemical analysis of EV release in R1441G LRRK2 cells showed similar behavior as in WT MEFs upon treatment with these agents (Figure 6A and Supplemental Figure 3). GW4869 inhibition of EV release increased cellular BMP content (Figure 6D), while enhanced EV release upon B-A1 treatment diminished intracellular total di-22:6-BMP and di-18:1-BMP (Figure 6E). Altogether, these results strongly support the notion that BMP is released by EV-mediated MVE exocytosis.

**Figure 6.**
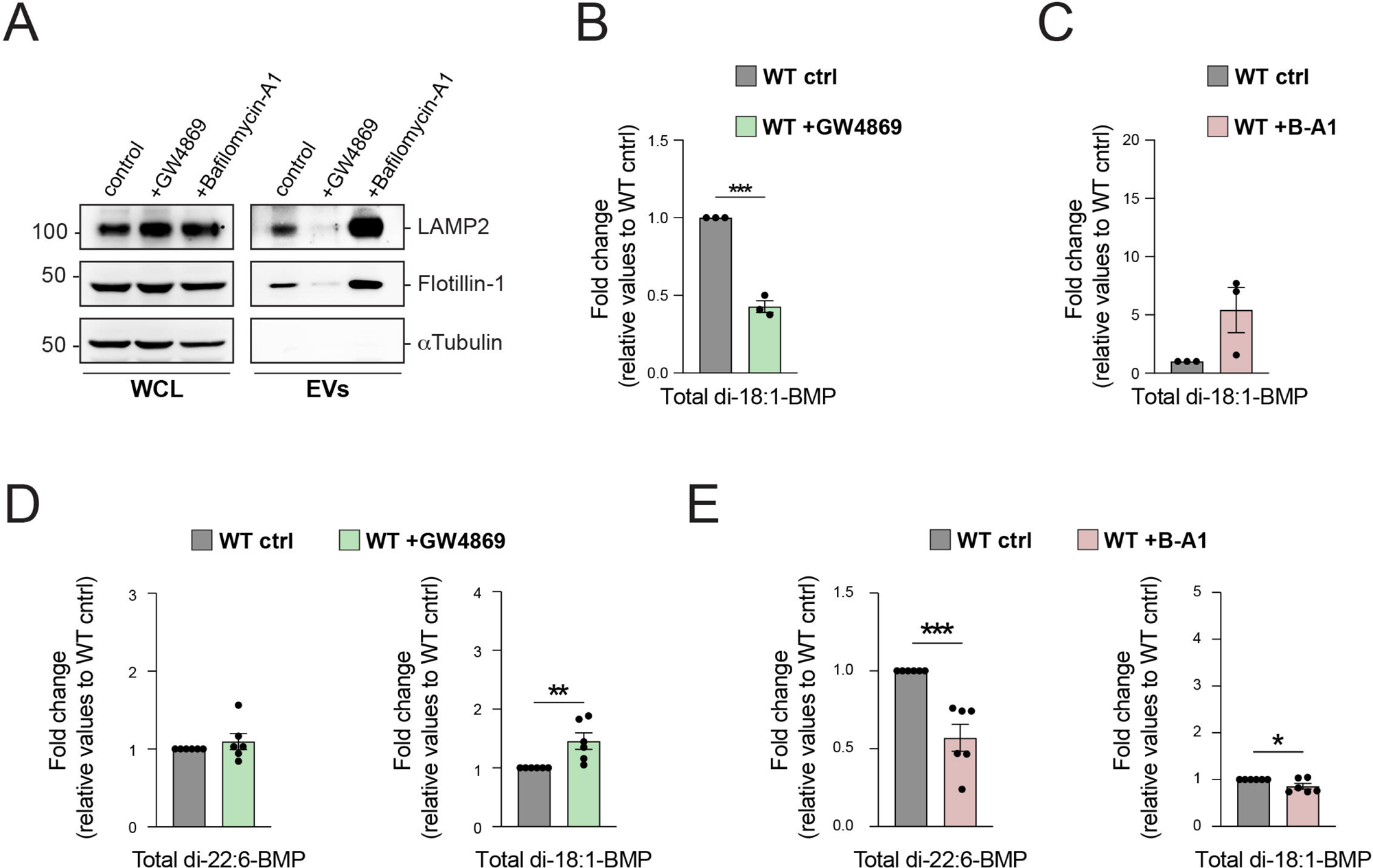
Pharmacological modulation of EV-mediated BMP exocytosis. **(A)** Whole cell lysates (WCL) and isolated EVs from WT MEF cells treated with 10µM GW4869 or 10nM bafilomycin-A1 (B-A1) for 24h were analyzed by immunoblotting. Representative immunoblots of LAMP2, Flotillin-1 and α-Tubulin are shown. Molecular weight marker mobility is shown in kDa. **(B-C)** UPLC-MS/MS determination of BMP isoforms normalized to protein content from WT MEF cells treated with 10nM B-A1 **(B)** or 10µM GW4869 **(C)** for 24h. Data shown as fold change relative to WT control MEF cells. Only BMP isoforms that were detected are shown. Data from 6 independent experiments (mean ± SEM). Significance determined by two-tailed unpaired *t* test *p<0.05, **p<0.01, ***p<0.001, ****p<0.0001. **(D-E)** Quantitation of BMP isoforms normalized to protein content in EVs isolated from MEF WT cells treated with 10nM B-A1 **(D)** or 10µM GW4869 **(E)** for 24h. Data shown as fold change relative to WT control MEF cells. Only BMP isoforms that were detected are shown. Data from 3 independent experiments (mean ± SEM). Significance determined by two-tailed unpaired *t* test *p<0.05, **p<0.01, ***p<0.001.

### Monitoring in vivo endolysosomal exocytosis and EV release in patient-derived G2019S LRRK2 fibroblasts

All our previous experiments investigating the role of LRRK2 in EV release regulation were conducted using mouse-derived cells. To further validate these findings, we employed skin fibroblasts derived from healthy donors (control) and G2019S LRRK2 PD patients. We first performed an immunofluorescence and biochemical analysis of the BMP-positive endolysosomal compartment in three independent skin fibroblast cell lines from each cohort. Consistent with our observations in MEFs, we observed a decrease in BMP immunostaining in LAMP1-positive endolysosomes of G2019S LRRK2 cells compared to control (Figure 7A and B). LAMP1 levels remained unchanged between the two cohorts (Figure 7A and C).

**Figure 7.**
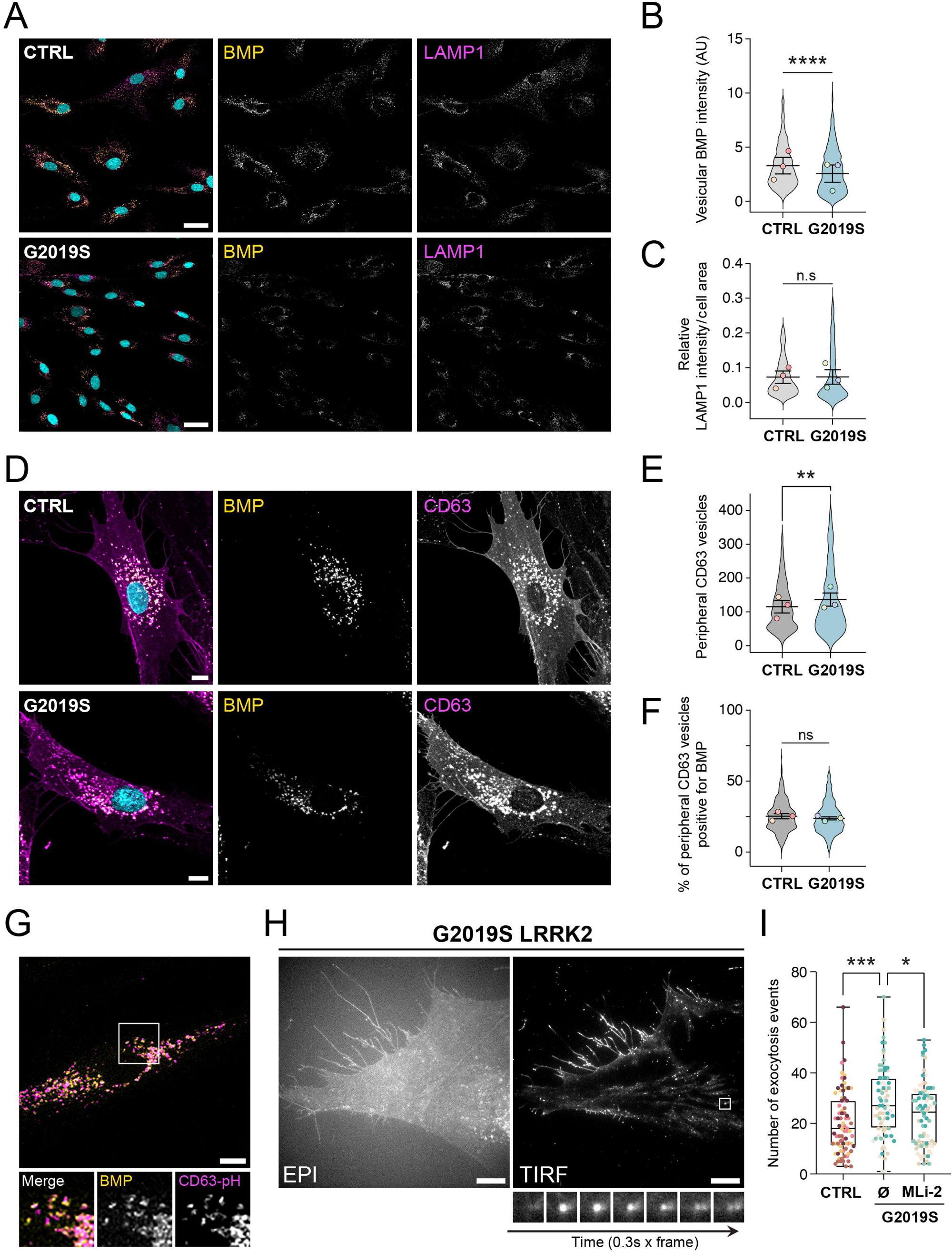
Patient-derived G2019S LRRK2 fibroblasts exhibit alterations of antibody-accessible BMP and increased endolysosomal exocytosis. (A) Confocal microscopy of endogenous BMP (yellow) and LAMP1 (magenta) immunofluorescence in control (CTRL) and LRRK2-G2019S mutant–derived fibroblasts. Scale bar: 40 µm. (B–C) Quantification of vesicular BMP intensity (B) and LAMP1 relative intensity (C) per cell area. Colored dots represent the mean value of three independent experiments (n=3 CTRL and n=3 G2019S LRRK2 fibroblast cell lines); violin plots show the distribution of individual cell data (60 cells per independent experiment). Significance determined by two-tailed paired *t* test; ****p < 0.0001. (D) Confocal microscopy of endogenous BMP (yellow) and CD63 (magenta) immunofluorescence in CTRL and G2019S LRRK2 fibroblasts. Scale bar: 10 µm. (E–F) Quantification of the number of CD63 vesicles in the peripheral cell region (E) and the percentage of peripheral CD63 vesicles positive for BMP (F). Colored dots represent mean values of three independent experiments (n=3 CTRL and n=3 G2019S LRRK2 fibroblast cell lines); violin plots show the distribution of individual cell data (60 cells per independent experiment). Significance determined by two-tailed paired *t* test; **p < 0.01. (G) A representative confocal microscopy image of endogenous BMP (yellow) and transduced CD63-pHluorin (magenta) in a CTRL human fibroblast cell line used for total internal reflection fluorescence (TIRF) microscopy experiments. Inset image shows co-localization between BMP and CD63-pHluorin in vesicular structures (a similar degree of co-localization was observed in G2019S LRRK2 cells; data not shown). Scale bar: 10 µm. (H) Representative epifluorescence (EPI) and TIRF microscopy images of the same cell from a G2019S LRRK2 patient-derived fibroblast cell line treated with vehicle (DMSO) overnight. Inset images shows a TIRF microscopy time-lapse sequence (0.3 s per frame) from a single pHluorin-CD63–positive fusion event at the plasma membrane. Scale bar: 10 µm. (I) Quantification of pHluorin-CD63–positive fusion events in stably expressing control (CTRL) and G2019S LRRK2 fibroblasts treated with vehicle (Ø) or 200 nM MLi-2 for 16 h. Each dot represents one cell (n=20 cells quantified from four CTRL and four G2019S LRRK2 cell lines). Significance determined by Tukey’s multiple comparisons test, ordinary one-way ANOVA; *p < 0.05, **p < 0.001.

Next, we biochemically characterized endolysosomes isolated by subcellular fractionation on discontinuous sucrose gradients. Interestingly, and consistent with previous studies reporting association of LRRK2 with isolated endolysosomes^48^, a pool of LRRK2 was detected in fractions positive for CLN5 and the endolysosomal marker CD63, but negative for tubulin (Supplemental Figure 4A). As seen in our whole cell lysate analysis (Figure 5C), relative CLN5 levels appeared increased in G2019S LRRK2 fibroblasts (Supplemental Figure 4A, bottom fractions). However, the relative amount of CLN5 co-peaking with isolated CD63-positive endolysosomes was comparable to that in control cells (Supplemental Figure 4A, endolysosome fractions). These results suggests that, in these cells, a considerable pool of newly synthesized CLN5 may not have reached its final destination at the endolysosome and may instead be retained in the ER. Such a delay in CLN5 trafficking could explain why, despite the apparent increase in CLN5 protein levels also found in mutant LRRK2 MEFs, the BMP biosynthetic rate did not differ from that observed in WT MEFs. Despite these changes in CLN5 expression, no statistically significant differences were detected in the total protein levels of CD63 between control and G2019S LRRK2 human fibroblasts (Supplemental Figure 4B). Collectively, these data indicate that human cells carrying pathogenic LRRK2 mutations exhibit alterations in antibody-accessible BMP comparable to those observed in MEFs, and that a pool of LRRK2 may be present in CD63/CLN5-positive endolysosomes likely enriched in BMP.

CD63-positive compartments have been implicated in exocytic processes rather than catabolic activities^49–51^. In line with these observations, our data indicating that a pool of LRRK2 co-fractionates with isolated CD63-positive endolysosomes prompted us to further examine the intracellular distribution of CD63 in these cells. An increased number of CD63-positive vesicles near the plasma membrane was quantified in mutant LRRK2 fibroblasts compared to control cells (Figure 7D and E). On average, ∼25% of CD63-positive peripheral endolysosomes were also positive for BMP, with no observable differences between control and mutant LRRK2 cells (Figure 7D and F). The peripheral enrichment observed in G2019S LRRK2 fibroblasts suggests enhanced recruitment of CD63-positive compartments toward exocytic sites at the cell surface, supporting the notion that these vesicles may eventually fuse with the plasma membrane to release EVs, as previously proposed^49,50^.

To determine more precisely whether LRRK2 kinase activity modulates the exocytosis of CD63-positive endolysosomes, we performed live-cell total internal reflection fluorescence (TIRF) microscopy using CD63-pHluorin, a genetically encoded fluorescent reporter of exosome release^47,50^. For these experiments, four independent control and G2019S LRRK2 patient-derived fibroblast cell lines were transduced to stably express CD63-pHluorin. The presence of BMP in CD63-pHluorin–positive endolysosomes was confirmed by immunofluorescence analysis after selection of transduced cells (Figure 7G). Consistent with recent findings in human induced pluripotent stem cell (iPSC)-derived neurons harboring the R1441H LRRK2 pathogenic mutation^52^, TIRF microscopy revealed an overall increase in endolysosomal exocytosis in the four G2019S LRRK2 fibroblast cell lines tested, which was significantly reduced upon MLi-2 treatment (Figure 7H and I; Videos 1 and 2). Together, these findings are consistent with our previous observations in MEFs (Figures 2E and 3E) and support our initial hypothesis that LRRK2 kinase activity drives exocytosis and release of BMP-enriched EVs.

## DISCUSSSION

Here we have shown that: (a) hyperactive LRRK2 kinase upregulates mass spectrometry-determined BMP levels and the extracellular vesicles (EVs) that release BMP in MEF cells. (b) Pharmacological inhibition of LRRK2 and GCase using MLi-2 and CBE, respectively, modulates BMP abundance in cells and isolated exosomal fractions in opposite directions without interfering with the kinetics of BMP biosynthetic or catabolic rates, although the effect of GCase inhibition on EV-associated BMP showed more variability across experiments. While PD-associated mutations in *LRRK2* increase kinase activity, *GBA1* pathogenic variants are associated with decreased GCase enzymatic function. Therefore, MLi-2 treatment represents a potential therapeutic rescue, whereas CBE phenocopies pathogenic conditions. (c) Although the hyperactive kinase mutation is associated with an overall increase in the BMP synthesis, neither MLi-2 or CBE affected BMP biosynthetic rate in the MEFs. (d) The expression of CLN5, a BMP synthase^45,46^, is upregulated in mouse and patient-derived fibroblasts with PD-associated LRRK2 mutations, consistent with the lipidomics data indicating elevated amounts of cellular BMP. (e) BMP release was modulated by pharmacological agents known to modulate EV secretion. (f) Finally, G2019S LRRK2 human fibroblasts exhibit enhanced endolysosomal exocytosis and EV release which decreased upon MLi-2 treatment. Together, these data indicate that the previously reported increase in urinary BMP levels in LRRK2 mutation carriers reflects dysregulated exocytosis of BMP-containing EVs.

Pioneering studies by Gruenberg and colleagues underscore the importance of BMP in endolysosome homeostastic regulation and generation of endolysosomal ILVs^14,18,19^ that become exosomes upon release. Subsequent studies revealed that cellular levels of this atypical phospholipid are invariably altered in many neurodegenerative disorders characterized by endolysosome dysfunction^24^. Recently, key lysosomal enzymes involved in distinct steps of the BMP biosynthetic pathway were identified, including CLN5 which plays a critical role in this process^45,46^. Our microscopy analysis revealed decreased antibody-accessible BMP levels in R1441G LRRK2 cells. In contrast, our targeted lipid pathway analysis measurements consistently showed higher total BMP levels in R1441G LRRK2 cells compared with WT cells. As we have reported previously^40^, BMP antibody detects only a sub-pool of “accessible” BMP while mass spectrometry detects all pools. Alternately, not all BMP isoforms may be detected equally well. Given the essential roles of BMP in endolysosomal catabolism, our immunofluorescence data predict that LRRK2 mutant cells have defective degradative capacity, consistent with recent reports^31,53^. In addition, in both MEFs and patient-derived mutant LRRK2 fibroblasts we found upregulation of the BMP-synthesizing enzyme CLN5 and, in MEFs, a concomitant increase in LAMP2 protein levels. These findings suggest that LRRK2 may participate in endolysosome biogenesis regulation. Indeed, in macrophages and microglia, LRRK2 regulates the levels of multiple lysosomal proteins by inhibiting TFEB and MiTF^53^, transcription factors that activate lysosome-related gene expression by binding to coordinated lysosomal expression and regulation (CLEAR) elements^54^. Although the promoter region of CLN5 contains a potential TFEB binding site^54^, formal evidence for TFEB-mediated transcription of CLN5 is still lacking. The increase in CLN5 (and LAMP2 in MEFs) protein expression may also reflect lysosomal stress^54^. Interestingly, MLi2 restored CLN5 levels in both murine-derived and human fibroblasts, further providing support that BMP is a pharmacodynamic biomarker of not just target modulation but also a potential disease-relevant lysosomal pathway modulation. Our findings of CLN5 upregulation in mutant LRRK2 MEFs, validated by similar results in patient-derived fibroblasts, further establish MEFs as a relevant cellular model system for studying LRRK2 kinase activity and its therapeutic targeting^55–58^. Taken together, our data indicate that LRRK2 activity modulates specific components of the lysosomal network, including CLN5 and LAMP2. Nevertheless, quantitative analysis of additional endolysosomal markers (LAMP1 and CD63) in human fibroblasts did not reveal statistically significant differences between control and mutant LRRK2 cells. The elevated LAMP2 expression observed in the engineered MEF clone expressing R1441G may reflect a cell type-specific effect, potentially linked to differential penetrance of LRRK2 signaling on the lysosome biogenesis response.

We and others reported that BMP is aberrantly high in urine derived from LRRK2 and GBA mutation-carriers^27–29^. These results have also been reproduced in animal models, in which urinary BMP levels decrease upon administration of MLi-2 and other LRRK2 kinase inhibitors^41–43^. The present study complements these previous observations by providing evidence that EV-mediated BMP release can be regulated by LRRK2 and, although less consistently, by GCase enzymatic activity. Consistent with this model, recent studies have shown that aberrant LRRK2 and GCase activities influence EV secretion in cellular and animal models of PD, including human patients^59–61^. Moreover, our data showing reduced exocytosis and EV release in mutant LRRK2 MEFs and patient-derived fibroblasts treated with MLi-2 is consistent with previous observations that LRRK2 kinase inhibition leads to surfactant accumulation in type 2 pneumocytes in the lung, likely as a consequence of impaired lysosome-related organelle exocytosis^42^. Moreover, a recent study also reporting decreased exocytosis of CD63-pHluorin–positive compartments in MLi-2–treated primary murine neurons and human induced pluripotent stem cell (iPSC)-derived neurons harboring pathogenic LRRK2 mutations^52^, further supports our conclusions.

BMP is enriched in urinary EVs^62^. However, other studies did not detect BMP enrichment in isolated EV fractions and concluded that BMP may be present in ILVs of endolysosomal subpopulations devoted to degradative activities rather than exocytic events^63,64^, as previously proposed^65^. Using our UPLC-MS/MS methodology, extracellular BMP can be efficiently detected in the urine of healthy and mutant LRRK2/GCase carriers^27–29^, and additionally, in EV fractions derived from both WT and R1441G LRRK2 MEFs (this study). How does LRRK2 influence release of BMP-positive EVs (BMP-EVs)? A subset of Rab proteins that are master regulators of intracellular membrane trafficking pathways are important LRRK2 substrates^5^. Some of these Rabs include Rab10, Rab12 and Rab35, and have been shown to play regulatory roles in endolysosomal exocytosis and EV release^66,67^. We thus speculate that phosphorylation of one or several of these Rabs may lead to enhanced BMP release. Indeed, G2019S LRRK2-mediated Rab35 phosphorylation has been proposed to promote EV-mediated α-synuclein release and propagation between cells^68,69^. It is likely that aberrant LRRK2 and GCase activities trigger exocytosis, and associated release of BMP-containing EVs, via a so-called clearance pathway activated due to accumulation of cytotoxic endolysosomal substrates^70,71^. The contribution of these phosphorylated Rabs to EV-mediated BMP release will be of interest for future work.

In summary, this work provides evidence that pathological LRRK2 activity, and to a more variable degree GCase dysfunction, enhance EV-mediated BMP release without altering its metabolism in cells. The observed changes induced by MLi-2 highlight the potential of LRRK2 inhibition as a therapeutic strategy to restore endolysosomal homeostasis by reducing BMP production and aberrant EV release. What is the clinical significance of elevated urinary BMP levels in PD patients? Inhibition of LRRK2 improves endolysosomal function^43^, consistent with increased LRRK2 and decreased GCase activities associated with pathogenic variants which worsen endolysosomal homeostasis^6,13^; this may eventually lead to endolysosomal exocytosis followed by EV-mediate BMP release. We speculate that BMP-EVs may harbor a distinct molecular repertoire that could perhaps inform more precisely disease progression or pathobiology. Future investigations will be needed to study the contribution of dysfunctional LRRK2 and GCase in exocytosis of specific BMP-positive endolysosomal subpopulations and to further characterize the role of BMP-EVs in the context of disease pathophysiology but also as a PD diagnostic tool.

## ACKNOWLEDGMENTS

This research was funded by a grant to K.M and A.L from The Michael J. Fox Foundation for Parkinson’s Research (MJFF-019043).

## AUTHOR CONTRIBUTIONS

K.M.M. and A.L. conceived the project with input from F.H. All experiments were carried out by E.M-S. and A.L. except for mass spectrometric analysis of BMP (F.H), and CLN5 biochemical analysis in human fibroblasts (M.C.). R.FS., M.E., A.G and M.J.M provided patient-derived fibroblasts. C.E. captured and analyzed transmission electron microscopy images. K.M., A.L., S.R.P. and F.H. oversaw the project. K.M. and A.L. obtained research funding. A.L. wrote and edited the manuscript with critical revision from all co-authors.

## COMPETING INTERESTS

Following authors declare no competing non-financial interests but the following competing financial interests: Kalpana M. Merchant, PhD, has consulted for AcureX, Caraway, Nitrase, Nura Bio, Retromer Therapeutics, Sinopia Biosciences, Vanqua, Vida Ventures and the Michael J. Fox Foundation for Parkinson’s Research. She has received research funding from the Michael J. Fox Foundation for Parkinson’s Research. Albert Lu, MD, PhD, and Suzanne R. Pfeffer, Ph.D. have received research funding from the Michael J. Fox Foundation for Parkinson’s Research. Frank Hsieh, PhD and Ms. Marianna Arnold are employed by Nextcea, Inc., which holds patent rights to the di-22:6-BMP and 2,2’-di-22:6-BMP biomarkers for neurological diseases involving lysosomal dysfunction (US 8,313,949, Japan 5,702,363, and Europe EP2419742).

## RELEVANT CONFLICTS OF INTEREST/FINANCIAL DISCLOSURES

Dr. Hsieh and Ms. Arnold are employed by Nextcea, Inc., which holds patent rights to the di-22:6-BMP and 2,20-di-22:6-BMP biomarkers for neurological diseases involving lysosomal dysfunction (US 8,313,949, Japan 5,702,363, and Europe EP2419742).

## FUNDING AGENCIES

This research was funded by the Michael J. Fox Foundation (MJFF-019043; MJFF-023914).

## ABBREVIATIONS

BMP, bis(monoacyglycerol)phosphate; CBE, conduritol β-epoxide; DHA, docosahexaenoic acid; GCase, glucocerebrosidase; ILV, intralumenal vesicle; LBPA, Lysobisphosphatidic acid; LRRK2, leucine-rich repeat kinase 2; PD, Parkinson’s disease; WT, wild type

## MATERIALS AND METHODS

### Cell culture, antibodies and other reagents

Human-derived skin fibroblasts^74^, wild type (WT) and R1441G LRRK2 mutant MEF cells were grown in Dulbecco’s modified Eagle’s media (DMEM) containing 10% fetal bovine serum, 2 mM L-glutamine, and penicillin (100 U/ ml)/streptomycin (100 mg/ml). Cell lines were cultured at 37°C with 5% CO2. Extracellular vesicle-free media was prepared by overnight ultracentrifugation of DMEM or RPMI supplemented with 10% FBS at 100,000 X g in a SW32Ti rotor. For bafilomycin-A1 (Sigma-Aldrich; catalog no. B1793) and GW4869 (SelleckChem; catalog no. S7609), the drug was added to cell media at 10 nM and 10 µM respectively and cells were cultured for 24 h before exosome collection. For MLi-2 (Tocris Bioscience; catalog no. 5756) and Conduritol β-Epoxide (MERK; catalog no. 234599) were added to cell media at 200nM and 300µM, respectively. In our experiments, 48-hour incubations were necessary to sustain full LRRK2/GCase inhibition throughout the EV collection period. EV biogenesis, BMP biosynthesis, and packaging into EVs are time-dependent processes; therefore, extended incubation and collection periods (≥48 h) were required to allow downstream effects of LRRK2/GCase inhibition on BMP production and release to manifest, and to obtain sufficient EV material for biochemical and lipidomic analyses. 0.167mM Oleic acid (Cambridge Isotope Laboratories; catalog no. DLM-10012-0.001) and Docosahexaenoic acid (Cambridge Isotope Laboratories; catalog no. DLM-10012-0.001) were conjugated with 0.0278mM fatty acid-free BSA (Sigma-Aldrich; A8806) at a 1:6 molar ratio. Primary antibodies diluted in PBS with 1% BSA (for immunofluorescence) or 5% skim milk (for immunoblotting) were mouse anti-lysobisphosphatidic acid (anti-BMP) clone 6C4 1:1000 (EMD Millipore; catalog no. MABT837), rat monoclonal anti-mouse LAMP2 1:1000 (Developmental Studies Hybridoma Bank, catalog no.GL2A7), mouse anti-Flotillin-1 1:1000 (BD Biosciences; catalog no.610821), rabbit anti-phospho-Rab10 1:1000 (Abcam; catalog no. ab230261), mouse anti-alpha tubulin 1:10000 (SigmaAldrich; catalog no. T5168), rabbit anti-CLN5 1:000 (Abcam; catalog no. ab170899), rabbit anti-phosphoLRRK2 1:000 (Abcam; catalog no. ab133450), rabbit anti-LAMP1 XP D2D11 1:1000 (Cell Signalling; catalog no.9091), rabbit anti-CD63 [EPR22458-280] 1:500 (Abcam; catalog no.AB252919), rabbit anti-LRRK2 1:500 (Abcam; catalog. No. AB133474), mouse anti-calnexin (BD Transduction; 610523) 1:1000 and rabbit anti-GFP (Abcam, catalog no. AB290) 1:500.

### Immunofluorescence

Wild type (WT) and R1441G LRRK2 mutant MEF cells were fixed with 3.7% (v/v) paraformaldehyde for 15 min, washed and permeabilized/blocked with 0.1% saponin/1% BSA-PBS before staining with anti-LAMP2 and anti-BMP antibodies followed by goat anti-rat AlexaFluor555 and donkey anti-mouse AlexaFluor488. Nuclei were stained using 0.1 mg/ml DAPI (SigmaAldrich; catalog no. D9542). Coverslips were mounted on glass slides with Mowiol. Microscopy images were acquired using a Zeiss LSM880 laser scanning spectral confocal microscope (Carl Zeiss) equipped with an Axio Observer 7 inverted microscope, a blue diode (405 nm), argon (488 nm), diode-pumped solid-state (561 nm), and HeNe (633 nm) lasers, and a Plan Apochromat 63×/NA1.4 oil-immersion objective lens. DAPI, Alexa Fluor 488, Alexa Fluor 555 were acquired sequentially using 405-, 488- and 561-nm laser lines; Acousto-Optical Beam Splitter as a beam splitter; and emission detection ranges 415-480 nm, 500-550 nm, 571-625 nm, respectively. The confocal pinhole was set at 1 Airy unit. All images were acquired in a 1,024 × 1,024-pixel format. BMP and LAMP2 intensities were quantified using CellProfiler^75^. All images for BMP-LAMP1 and BMP-CD63 analyses in human fibroblasts were acquired in tile-scan mode (5 × 5 field of view) with a z-step of 5, yielding a 2354 × 2354-pixel format. Tile-scan images were stitched with a 10% overlap, and maximum-intensity projections were generated using the Zeiss Zen Blue software. BMP and LAMP1 fluorescence intensities, as well as the number of peripheral CD63-positive vesicles and the percentage of peripheral CD63 vesicles positive for BMP, were quantified using Fiji (ImageJ) software.

### Lentivirus production and cell transduction

To produce CD63-pHluorin lentivirus, a lentivirus vector containing the CD63-pHluorin–expressing cassette^47^ was combined with third-generation lentiviral packaging mix (1:1:1 mix of pMD2.G, pRSV-Rev, and pMDLg/pRRE) at a 1:1 ratio and transfected into HEK293T cells using Polyethylenimine (PEI, #23966-2, PolySciences Inc.), as previously described^76^. Transfected cells were grown for 72 hr and then the supernatant containing the viral particles was collected and passed through a sterile 0.45 μm Millex syringe filter (Millipore). Human-derived skin fibroblasts grown in 60 mm culture dishes were infected by adding a 1:1 mixture of complete DMEM medium and viral supernatant supplemented with 8 mg/mL polybrene, followed by a 24h incubation. The supernatant was removed, and cells were grown in fresh DMEM medium for 48-72 hr. Infected cells were selected by growing in DMEM medium supplemented with 1-2 mg/mL puromycin (Sigma- Aldrich) for 3-5 days. Cells stably expressing CD63-pHluorin were recovered in complete DMEM medium lacking puromycin for 48-72 hours.

### Total Internal Reflection Fluorescence (TIRF) Live Microscopy

Control and G2019S LRRK2 human fibroblasts stably expressing pHluorin-CD63 were cultured on 25 mm coverslips and treated with DMSO (vehicle) or 200 nM MLi-2. After 16 h, cells were imaged using a Zeiss LSM880 Airyscan Elyra PS.1 laser scanning spectral confocal microscope (Carl Zeiss) equipped with an Axio Observer Andor1 inverted microscope. Imaging was performed with an α Plan-Apochromat TIRF 100×/1.46 NA oil DIC objective, a BP495–575 + LP750 filter set, and an sCMOS camera operated in TIRF mode. Excitation was achieved with an HR diode laser (488 nm, 200 mW). Time-lapse videos were acquired at 0.30 s per frame for a total of 540 frames per cell analyzed. All imaging experiments were performed at 37°C in a humidified incubator (5% CO₂). Fusion events for each cell were detected and quantified as sudden increases in fluorescence at the cell surface using the AMvBE macro in Fiji (ImageJ), as described^77^. Twenty cells were imaged and quantified for each of the four fibroblast cell lines within each cohort.

### Immunoblotting

Cells were lysed in lysis buffer (50 mM Hepes, 150 mM KCl, 1% Triton X-100, 5 mM MgCl2, pH 7.4) supplemented with a protease/ phosphatase inhibitor cocktail (1mMNa3VO4, 10mMNaF, 1 mM PMSF, 10 μg/ml leupeptin, and 10 μg/ml aprotinin). After centrifugation at 12,000 g for 10 min, protein concentrations were measured in the cleared lysates. Equal protein amounts were loaded in lanes corresponding to whole-cell lysate (WCL) fractions and normalized based on α- Tubulin levels. For the extracellular vesicle (EV) fractions, all protein recovered from EV pellets after isolation was loaded. Exosome final pellets were resuspended in SDS-PAGE loading dye (50 mM Tris- HCl pH 6.8, 6% glycerol, 2% SDS, 0.2% bromophenol blue), heated briefly at 95°C, resolved by 10% SDS-PAGE gel and transferred onto nitrocellulose membranes (Bio-Rad Laboratories; catalog no. 1620115) using a Bio-Rad Trans-Blot system. Membranes were blocked with 5% skim milk in TBS withTween-20 for 60 min at RT. Primary antibodies were diluted in blocking buffer and incubated overnight at 4°C. HRP-conjugated secondary antibodies (Bio-Rad Laboratories) diluted in blocking buffer at 1:3000 were incubated for 60 min at RT and developed using EZ-ECL (Biological Industries) or SuperSignal West Femto (Thermo Scientific; catalog no. 34094). Blots were imaged using an ImageQuant LAS 4000 system (GE Healthcare) and quantified using ImageJ software.

### Transmission Electron Microscopy

Cells in culture were washed in PBS and fixed for 1h in 2.5% glutaraldehyde in 0.1 M phosphate buffer (PB) at RT. Next, samples were slowly and gently scraped and pelleted in 1.5 ml tubes. Pellets were washed in PB and incubated with 1% OsO4 for 90 min at 4 °C. Then samples were dehydrated, embedded in Spurr and sectioned using Leica ultramicrotome (Leica Microsystems). Ultrathin sections (50-70 nm) were stained with 2% uranyl acetate for 10 min, a lead-staining solution for 5 min and observed using a transmission electron microscope, JEOL JEM- 1010 fitted with a Gatan Orius SC1000 (model 832) digital camera. Multivesicular Endosomes (MVE) and Intralumenal Vesicles (ILV) were identified by morphology and MVE area and ILV number were measured using ImageJ.

### Extracellular vesicle isolation

MEF adherent cell cultures were seeded at 3-4*10^6^ cells/plate in EV- free complete DMEM and grown for 24-48 h. Cell cultures were centrifuged at 300 g, 5 min to pellet cells, and supernatants were centrifuged again at 2100 g, 20 min to pellet dead cells. After this, filtration through a 0.22 μm filter unit (Millipore) was performed. Filtered media were ultracentrifuged for 75 min at 100,000 g in SW32Ti rotor to pellet extracellular vesicles. The supernatant was discarded, and the pellet (small EVs) re-suspended in phosphate-buffered saline (PBS) and centrifuged again for 75 min at 100,000 g in 140AT rotor. exosome pellets were frozen and stored at -80°C prior to further processing. In all EV-related experiments, we seeded the same number of EV-producing cells per condition, and the resulting EV-derived data (from both immunoblotting and lipidomics analyses) were normalized to the corresponding whole cell lysate (WCL) protein content to ensure comparability across conditions.

### Lipidomic analysis

Targeted quantitative ultra-performance liquid chromatography tandem mass spectrometry was used to accurately quantitate the three geometrical isomers (2,2-, 2,3- and 3,3-) of di-22:6-BMP and di-18:1-BMP in control or treated MEF cells. Lipidomic analyses were conducted by Nextcea, Inc. as previously described^66^. Standard curves were prepared using authentic BMP reference standards. Protein was determined by bicinchoninic acid protein assay. A multiplexed quantitative ultraperformance liquid chromatography tandem mass spectrometry method was used to simultaneously quantitate sample glucosylceramides (GluCer d18:1/16:0, d18:1/18:0, d18:1/22:0, d18:1/24:0, d18:1/24:1), galactosylceramides (GalCer d18:1/16:0, d18:1/18:0, d18:1/22:0, d18:1/24:0, d18:1/24:1), glucosylsphingosine (GluSph 18:1), and galactosylsphingosine (GalSph 18:1). Standard curves were prepared from related standards using a class-based approach. Internal standards were used for each analyte reported. A SCIEX Triple Quad 7500 mass spectrometer was used in positive electrospray ionization mode for detection (SCIEX, Framingham, MA, USA). Injections were made using a Shimadzu Nexera XR UPLC (ultraperformance liquid chromatography) system (Shimadzu Scientific Instruments, Kyoto, Japan). The instruments were controlled by SCIEX OS 2.0 software.

### Metabolic labeling

WT and R1441G LRRK2 mutant MEF cells were treated with 200nM MLi-2 or 300µM CBE for 16h. Cells were washed twice with PBS and incubated in serum-free DMEM ± MLi-2 or CBE for 1h. Next, Incubate cells in 0.167mM 13C-labeled Oleic Acid (OA) and 0.167mM deuterated docosahexaenoic acid (DHA) in serum-free DMEM containing 0.0278mM BSA for 25 min. Cells were washed with serum-free DMEM and chased with serum-free DMEM for short time points (0, 15, 30, 45, 60min) or complete DMEM for long timepoints (8, 24 and 48h). Cells were washed with PBS, trypsinized and collected in cold PBS. Cells were pelleted by centrifugation at 1000rpm for 7min and stored at -80°C prior to further processing.

### GCase activity

GCase activity was measured by using the fluorogenic substrate PFB-FDGlu (Invitrogen, Carlsbad, CA). WT and R1441G LRRK2 mutant MEF cells were treated with 300μM CBE for 24h. After this, 150 μg/ml PFB-FDGlu (Thermo Fisher) was then added to cells followed by incubation for 2h at 37°C. Cells were then washed twice with PBS, trypsinized and fixed in cold 2% paraformaldehyde-PBS for 30 min. Cells were washed twice in PBS before assayed on a FACS analyzer (FACScan; BD) equipped with a 488 nm laser and 530 nm-FITC filter. Data were analyzed using Flowjo software (Tree Star).

### Subcellular fractionation

Endolysosomes were isolated by ultracentrifugation of discontinuous sucrose density gradients as previously described^78^. Briefly, control and G2019S LRRK2 fibroblasts were collected using a homogenization buffer containing 3mM Imidazole, 250nM sucrose, pH 7,4 and proteases inhibitors. Cell pellets were homogenized by 90 passages through a 27-gauge needle. Complete homogenization was assessed by phase-contrast microscopy. After this, samples were centrifuged for 10 min at 1000 g. The Post-nuclear supernatant (PNS) fractions were collected and adjusted to 40,2% (w/v) sucrose. PNS fractions were then loaded at the bottom of a micro- ultracentrifuge tube for Himac P70AT(RP70AT) Fixed Angle Rotor. Then, 35% sucrose, 25% sucrose, and 8% sucrose layers were poured stepwise over the PNS bottom layer. The discontinuous sucrose gradient was centrifuged for 3 h at 100,000 g using a Himac CS-(F)NX (Eppendorf) micro- ultracentrifuge. After this, 15 fractions were collected, and protein was trichloroacetic acid (TCA)- precipitated before SDS-PAGE analysis.

### Stabsbcs

Results are expressed as the mean ± SEM. Means were compared using Student’s t test when two experimental condiãons were independently compared. Unless otherwise noted, staãsãcal significance between mulãple comparisons was assessed via one-way or two-way ANOVA with uncorrected Fisher’s LSD test using GraphPad Prism version 9.

---

**Supplemental Figure 1.** Further characterization of MEF-derived EV fractions. (A-C) No significant differences in EV release between MLi-2/CBE-treated and untreated WT MEF cells. Quantification of LAMP2 and Flotillin-1 levels relative to WT control MEF cells in WCL **(A)** and isolated EVs **(B-C).** Data from 7-8 independent experiments (mean ± SEM). Significance determined by ordinary one-way ANOVA, uncorrected Fisher’s LSD. **(D)** and **(E)**, Characterization of WT or R1441G LRRK2 MEF-derived purified EVs by Nanoparticle Tracking Analysis (NTA). **(D)** Representative plots of EV size distribution in each indicated condition. **(E)** Yield comparison of MEF-derived EVs from each indicated condition determined by NTA. Each colored dot represents an independent experiment. **(F,G)** No significant differences in cellular BMP levels between MLi-2/CBE-treated and untreated WT MEFs, as measured by UPLC-MS/MS. **(H)** Table showing absolute values of UPLC-MS/MS measurements of cellular docosahexaenoyl (22:6)-, arachidonyl (20:4)-, oleoyl (18:1)-, and linoleyl (18:2)-BMP species from 3 representative independent experiments. **(I)** No significant differences in EV-associated BMP levels between MLi-2/CBE-treated and untreated WT MEF cells, as measured by UPLC-MS/MS. Lipidomics measurements of cellular **(F,G)** and EV-associated **(I)** BMP isoforms normalized to protein content from cells treated with 200nM MLi-2 or 300µM CBE for 48h, or left untreated (ctrl). Data shown as fold change relative to WT control MEF cells. Only BMP isoforms that were detected are shown. Data from 6-7 independent experiments (mean ± SEM). Significance determined by ordinary two-tailed paired *t* test; ns, not significant.

**Supplemental Figure 2.** **Quantitative analysis of GCase substrates in MEF cells and EV fractions. (A-B)** Lipidomic determination of Glucosylceramide (GlcCer), Galactosylceramide (GalCer) and Glucosylsphingosine (GlcSph) isoforms normalized to protein content from WT and R1441G LRRK2 mutant MEF cells **(A)** and isolated EVs **(B)** treated with 300µM CBE or 200nM MLi-2 for 48h. Heatmap showing values as fold change relative to R1441G LRRK2 mutant MEF cell control. Only GlcCer, GalCer and GlcSph isoforms that were detected are shown. Data from 3-9 independent experiments (mean ± SEM). Significance determined by ordinary two-way ANOVA, uncorrected Fisher’s LSD *p<0.05, **p<0.01, ***p<0.001 ****p<0.0001.

**Supplemental Figure 3.** P**h**armacological **modulation of EV-mediated BMP exocytosis in mutant LRRK2 MEF cells.** Whole cell lysates (WCL) and isolated EVs from R1441G LRRK2 MEF cells treated with 10µM GW4869 or 10nM bafilomycin-A1 for 24h were analyzed by immunoblotting. Representative images of LAMP2, Flotillin-1 and α-Tubulin levels are shown. Molecular weight marker mobility is shown in kDa.

**Supplemental Figure 4.** A**n**alysis **of endolysosomal fractions in control and G2019S LRRK2 human- derived skin fibroblasts.** (A) Immunoblots showing fractions recovered after ultacentrifugation of sucrose gradients of cell extracts from control (CTRL) and G2019S LRRK2 mutant-derived fibroblasts. Endolysosomes fractions correspond to fractions 8-10. Representative immunoblots for LRRK2, CLN5, CD63 and α-tubulin are shown. Plots on the right show quantification of LRRK2 and CLN5 band intensities. n=3 CTRL and n=3 G2019S LRRK2 fibroblast cell lines. Molecular weight marker mobility is shown in kDa. (B) Whole cell lysates from CTRL individuals and G2019S LRRK2 mutant-derived fibroblasts analyzed by immunoblotting. Representative blots of CD63 and calnexin are shown. Plot on the right shows quantification of relative CD63 protein levels in whole-cell lysates normalized to Calnexin (used as a loading control). Molecular weight marker mobility is shown in kDa. Significance determined by ordinary two-tailed paired *t* test; ns, not significant.

Video 1. **Time-lapse of vehicle (DMSO)-treated human G2019S LRRK2 fibroblast cell stably expressing CD63-pHluorin.** Timestamp on upper left corner indicates seconds. Fusion events are indicated by white circles. Scale bar: 10 µm. Images were acquired at ∼3.33 frames per sec.

Video 2. **Time-lapse of MLi-2–treated human G2019S LRRK2 fibroblast cell stably expressing CD63- pHluorin.** Timestamp on upper left corner indicates seconds. Fusion events are indicated by white circles. Scale bar: 10 µm. Images were acquired at ∼3.33 frames per sec.

## Notes

### Summary of Updates

In this revised version, we include new data obtained from mutant LRRK2 patient-derived fibroblasts, including real-time monitoring of extracellular vesicle (EV) release using live TIRF microscopy, new immunofluorescence analyses, and subcellular biochemical fractionation experiments. Accordingly, we have updated our manuscript's results, discussion and methods sections, and have incorporated additional references that support and contextualize our findings. Finally, we have softened our conclusions regarding the contribution of GCase to EV-mediated BMP secretion.

## REFERENCES

1. Smith LJ, Lee CY, Menozzi E, Schapira AHV. Genetic variations in GBA1 and LRRK2 genes: Biochemical and clinical consequences in Parkinson disease. Front Neurol. 2022;13:971252. doi:10.3389/fneur.2022.971252

2. Yue Z, Lachenmayer ML. Genetic LRRK2 models of Parkinson’s disease: Dissecting the pathogenic pathway and exploring clinical applications. Mov Disord. 2011;26(8):1386–1397. doi:10.1002/mds.23737

3. Pont-Sunyer C, Tolosa E, Caspell-Garcia C, et al. The prodromal phase of leucine-rich repeat kinase 2-associated Parkinson disease: Clinical and imaging Studies. Mov Disord. 2017;32(5):726–738. doi:10.1002/mds.26964

4. Fan Y, Nirujogi RS, Garrido A, et al. R1441G but not G2019S mutation enhances LRRK2 mediated Rab10 phosphorylation in human peripheral blood neutrophils. Acta Neuropathol. 2021;142(3):475–494. doi:10.1007/s00401-021-02325-z

5. Pfeffer SR. LRRK2 phosphorylation of Rab GTPases in Parkinson’s disease. FEBS Lett. 2022;10.1002/1873-3468.14492. doi:10.1002/1873-3468.14492

6. Roosen DA, Cookson MR. LRRK2 at the interface of autophagosomes, endosomes and lysosomes. Mol Neurodegener. 2016;11(1):73. doi:10.1186/s13024-016-0140-1

7. Sidransky E, Lopez G. The link between the GBA gene and parkinsonism. Lancet Neurol. 2012;11(11):986–998. doi:10.1016/S1474-4422(12)70190-4

8. Duran R, Mencacci NE, Angeli AV, et al. The glucocerobrosidase E326K variant predisposes to Parkinson’s disease, but does not cause Gaucher’s disease. Mov Disord. 2013;28(2):232–236. doi:10.1002/mds.25248

9. Fredriksen K, Aivazidis S, Sharma K, et al. Pathological α-syn aggregation is mediated by glycosphingolipid chain length and the physiological state of α-syn in vivo. Proc Natl Acad Sci U S A. 2021;118(50):e2108489118. doi:10.1073/pnas.2108489118

10. Stojkovska I, Wani WY, Zunke F, et al. Rescue of α-synuclein aggregation in Parkinson’s patient neurons by synergistic enhancement of ER proteostasis and protein trafficking. Neuron. 2022;110(3):436–451.e11. doi:10.1016/j.neuron.2021.10.032

11. Ysselstein D, Nguyen M, Young TJ, et al. LRRK2 kinase activity regulates lysosomal glucocerebrosidase in neurons derived from Parkinson’s disease patients. Nat Commun. 2019;10(1):5570. doi:10.1038/s41467-019-13413-w

12. Kedariti M, Frattini E, Baden P, et al. LRRK2 kinase activity regulates GCase level and enzymatic activity differently depending on cell type in Parkinson’s disease. NPJ Parkinsons Dis. 2022;8(1):92. doi:10.1038/s41531-022-00354-3

13. Do J, McKinney C, Sharma P, Sidransky E. Glucocerebrosidase and its relevance to Parkinson disease. Mol Neurodegener. 2019;14(1):36. doi:10.1186/s13024-019-0336-2

14. Gruenberg J. Life in the lumen: The multivesicular endosome. Traffic. 2020;21(1):76–93. doi:10.1111/tra.12715

15. Gallala, H. D., & Sandhoff, K. (2011). Biological function of the cellular lipid BMP-BMP as a key activator for cholesterol sorting and membrane digestion. Neurochemical research, 36(9), 1594–1600. 10.1007/s11064-010-0337-6

16. Alattia JR, Shaw JE, Yip CM, Privé GG. Molecular imaging of membrane interfaces reveals mode of beta-glucosidase activation by saposin C. Proc Natl Acad Sci U S A. 2007;104(44):17394–17399. doi:10.1073/pnas.0704998104

17. Abdul-Hammed M, Breiden B, Schwarzmann G, Sandhoff K. Lipids regulate the hydrolysis of membrane bound glucosylceramide by lysosomal β-glucocerebrosidase. J Lipid Res. 2017;58(3):563–577. doi:10.1194/jlr.M073510

18. Kobayashi T, Stang E, Fang KS, de Moerloose P, Parton RG, Gruenberg J. A lipid associated with the antiphospholipid syndrome regulates endosome structure and function. Nature. 1998;392(6672):193-197. doi:10.1038/32440

19. Matsuo H, Chevallier J, Mayran N, et al. Role of LBPA and Alix in multivesicular liposome formation and endosome organization. Science. 2004;303(5657):531-534. doi:10.1126/science.1092425

20. Mason, R. J., Stossel, T. P., & Vaughan, M. (1972). Lipids of alveolar macrophages, polymorphonuclear leukocytes, and their phagocytic vesicles. The Journal of clinical investigation, 51(9), 2399–2407. 10.1172/JCI107052

21. Huterer, S., & Wherrett, J. (1979). Metabolism of bis(monoacylglycero)phosphate in macrophages. Journal of lipid research, 20(8), 966–973.

22. Bouvier, J., Zemski Berry, K. A., Hullin-Matsuda, F., Makino, A., Michaud, S., Geloën, A., Murphy, R. C., Kobayashi, T., Lagarde, M., & Delton-Vandenbroucke, I. (2009). Selective decrease of bis(monoacylglycero)phosphate content in macrophages by high supplementation with docosahexaenoic acid. Journal of lipid research, 50(2), 243–255. 10.1194/jlr.M800300-JLR200

23. Grabner, G. F., Fawzy, N., Schreiber, R., Pusch, L. M., Bulfon, D., Koefeler, H., Eichmann, T. O., Lass, A., Schweiger, M., Marsche, G., Schoiswohl, G., Taschler, U., & Zimmermann, R. (2020). Metabolic regulation of the lysosomal cofactor bis(monoacylglycero)phosphate in mice. Journal of lipid research, 61(7), 995–1003. 10.1194/jlr.RA119000516

24. Akgoc Z, Sena-Esteves M, Martin DR, Han X, d’Azzo A, Seyfried TN. Bis(monoacylglycero)phosphate: a secondary storage lipid in the gangliosidoses. J Lipid Res. 2015;56(5):1006–1013. doi:10.1194/jlr.M057851

25. Boland S, Swarup S, Ambaw YA, et al. Deficiency of the frontotemporal dementia gene GRN results in gangliosidosis. Nat Commun. 2022;13(1):5924. doi:10.1038/s41467-022-33500-9

26. Goursot A, Mineva T, Bissig C, Gruenberg J, Salahub DR. Structure, dynamics, and energetics of lysobisphosphatidic acid (LBPA) isomers. J Phys Chem B. 2010;114(47):15712–15720. doi:10.1021/jp108361d

27. Alcalay RN, Hsieh F, Tengstrand E, et al. Higher Urine bis(Monoacylglycerol)Phosphate Levels in LRRK2 G2019S Mutation Carriers: Implications for Therapeutic Development. Mov Disord. 2020;35(1):134–141. doi:10.1002/mds.27818

28. Merchant KM, Simuni T, Fedler J, et al. LRRK2 and GBA1 variant carriers have higher urinary bis(monacylglycerol) phosphate concentrations in PPMI cohorts. NPJ Parkinsons Dis. 2023;9(1):30. doi:10.1038/s41531-023-00468-2

29. Gomes S, Garrido A, Tonelli F, et al. Elevated urine BMP phospholipids in LRRK2 and VPS35 mutation carriers with and without Parkinson’s disease. NPJ Parkinsons Dis. 2023;9(1):52. doi:10.1038/s41531-023-00482-4

30. Jennings D, Huntwork-Rodriguez S, Vissers MFJM, et al. LRRK2 Inhibition by BIIB122 in Healthy Participants and Patients with Parkinson’s Disease. Mov Disord. 2023;38(3):386–398. doi:10.1002/mds.29297

31. Henry AG, Aghamohammadzadeh S, Samaroo H, et al. Pathogenic LRRK2 mutations, through increased kinase activity, produce enlarged lysosomes with reduced degradative capacity and increase ATP13A2 expression. Hum Mol Genet. 2015;24(21):6013–6028. doi:10.1093/hmg/ddv314

32. Hockey LN, Kilpatrick BS, Eden ER, et al. Dysregulation of lysosomal morphology by pathogenic LRRK2 is corrected by TPC2 inhibition. J Cell Sci. 2015;128(2):232–238. doi:10.1242/jcs.164152

33. de Rus Jacquet A, Tancredi JL, Lemire AL, DeSantis MC, Li WP, O’Shea EK. The LRRK2 G2019S mutation alters astrocyte-to-neuron communication via extracellular vesicles and induces neuron atrophy in a human iPSC-derived model of Parkinson’s disease. Elife. 2021;10:e73062. doi:10.7554/eLife.73062

34. Albanese F, Mercatelli D, Finetti L, et al. Constitutive silencing of LRRK2 kinase activity leads to early glucocerebrosidase deregulation and late impairment of autophagy in vivo. Neurobiol Dis. 2021;159:105487. doi:10.1016/j.nbd.2021.105487

35. Kowal J, Arras G, Colombo M, et al. Proteomic comparison defines novel markers to characterize heterogeneous populations of extracellular vesicle subtypes. Proc Natl Acad Sci U S A. 2016;113(8):E968–E977. doi:10.1073/pnas.1521230113

36. Mathieu M, Névo N, Jouve M, et al. Specificities of exosome versus small ectosome secretion revealed by live intracellular tracking of CD63 and CD9. Nat Commun. 2021;12(1):4389. doi:10.1038/s41467-021-24384-2

37. Ferreira JV, da Rosa Soares A, Ramalho J, et al. LAMP2A regulates the loading of proteins into exosomes. Sci Adv. 2022;8(12):eabm1140. doi:10.1126/sciadv.abm1140

38. Gardiner C, Ferreira YJ, Dragovic RA, Redman CW, Sargent IL. Extracellular vesicle sizing and enumeration by nanoparticle tracking analysis. J Extracell Vesicles. 2013; 2: 10.3402/jev.v2i0.19671. doi:10.3402/jev.v2i0.19671

39. Bachurski D, Schuldner M, Nguyen PH, et al. Extracellular vesicle measurements with nanoparticle tracking analysis - An accuracy and repeatability comparison between NanoSight NS300 and ZetaView. J Extracell Vesicles. 2019;8(1):1596016. doi:10.1080/20013078.2019.1596016

40. Lu A, Hsieh F, Sharma BR, Vaughn SR, Enrich C, Pfeffer SR. CRISPR screens for lipid regulators reveal a role for ER-bound SNX13 in lysosomal cholesterol export. J Cell Biol. 2022;221(2):e202105060. doi:10.1083/jcb.202105060

41. Fuji RN, Flagella M, Baca M, et al. Effect of selective LRRK2 kinase inhibition on nonhuman primate lung. Sci Transl Med. 2015;7(273):273ra15. doi:10.1126/scitranslmed.aaa3634

42. Baptista MAS, Merchant K, Barrett T, et al. LRRK2 inhibitors induce reversible changes in nonhuman primate lungs without measurable pulmonary deficits. Sci Transl Med. 2020;12(540):eaav0820. doi:10.1126/scitranslmed.aav0820

43. Jennings D, Huntwork-Rodriguez S, Henry AG, et al. Preclinical and clinical evaluation of the LRRK2 inhibitor DNL201 for Parkinson’s disease. Sci Transl Med. 2022;14(648):eabj2658. doi:10.1126/scitranslmed.abj2658

44. Nyame K, Xiong J, Alsohybe HN, et al. PLA2G15 is a BMP hydrolase and its targeting ameliorates lysosomal disease. Nature. 2025;642(8067):474–483. doi:10.1038/s41586-025-08942-y

45. Medoh UN, Hims A, Chen JY, et al. The Batten disease gene product CLN5 is the lysosomal bis(monoacylglycero)phosphate synthase. Science. 2023;381(6663):1182–1189. doi:10.1126/science.adg9288

46. Bulfon D, Breithofer J, Grabner GF, et al. Functionally overlapping intra- and extralysosomal pathways promote bis(monoacylglycero)phosphate synthesis in mammalian cells. Nat Commun. 2024;15(1):9937. doi:10.1038/s41467-024-54213-1

47. Lu A, Wawro P, Morgens DW, Portela F, Bassik MC, Pfeffer SR. Genome-wide interrogation of extracellular vesicle biology using barcoded miRNAs. Elife. 2018;7:e41460. doi:10.7554/eLife.41460

48. Bentley-DeSousa A, Roczniak-Ferguson A, Ferguson SM. A STING-CASM-GABARAP pathway activates LRRK2 at lysosomes. J Cell Biol. 2025;224(2):e202310150. doi:10.1083/jcb.202310150

49. Mittelbrunn M, Gutiérrez-Vázquez C, Villarroya-Beltri C, et al. Unidirectional transfer of microRNA-loaded exosomes from T cells to antigen-presenting cells. Nat Commun. 2011;2:282. doi:10.1038/ncomms1285

50. Verweij FJ, Bebelman MP, Jimenez CR, et al. Quantifying exosome secretion from single cells reveals a modulatory role for GPCR signaling. J Cell Biol. 2018;217(3):1129–1142. doi:10.1083/jcb.201703206

51. Verweij FJ, Bebelman MP, George AE, et al. ER membrane contact sites support endosomal small GTPase conversion for exosome secretion. J Cell Biol. 2022;221(12):e202112032. doi:10.1083/jcb.202112032

52. Palumbos SD, Popolow J, Goldsmith J, Holzbaur ELF. Autophagic stress activates distinct compensatory secretory pathways in neurons. Proc Natl Acad Sci U S A. 2025;122(28):e2421886122. doi:10.1073/pnas.2421886122

53. Yadavalli N, Ferguson SM. LRRK2 suppresses lysosome degradative activity in macrophages and microglia through MiT-TFE transcription factor inhibition. Proc Natl Acad Sci U S A. 2023;120(31):e2303789120. doi:10.1073/pnas.2303789120

54. Sardiello M, Palmieri M, di Ronza A, et al. A gene network regulating lysosomal biogenesis and function. Science. 2009;325(5939):473-477. doi:10.1126/science.1174447

55. Dhekne HS, Yanatori I, Gomez RC, et al. A pathway for Parkinson’s Disease LRRK2 kinase to block primary cilia and Sonic hedgehog signaling in the brain. Elife. 2018;7:e40202. doi:10.7554/eLife.40202

56. Sobu Y, Wawro PS, Dhekne HS, Yeshaw WM, Pfeffer SR. Pathogenic LRRK2 regulates ciliation probability upstream of tau tubulin kinase 2 via Rab10 and RILPL1 proteins. Proc Natl Acad Sci U S A. 2021;118(10):e2005894118. doi:10.1073/pnas.2005894118

57. Dhekne HS, Yanatori I, Vides EG, et al. LRRK2-phosphorylated Rab10 sequesters Myosin Va with RILPL2 during ciliogenesis blockade. Life Sci Alliance. 2021;4(5):e202101050. doi:10.26508/lsa.202101050

58. Kania E, Long JS, McEwan DG, et al. LRRK2 phosphorylation status and kinase activity regulate (macro)autophagy in a Rab8a/Rab10-dependent manner. Cell Death Dis. 2023;14(7):436. doi:10.1038/s41419-023-05964-0

59. Papadopoulos VE, Nikolopoulou G, Antoniadou I, et al. Modulation of β-glucocerebrosidase increases α-synuclein secretion and exosome release in mouse models of Parkinson’s disease. Hum Mol Genet. 2018;27(10):1696–1710. doi:10.1093/hmg/ddy075

60. Cerri S, Ghezzi C, Ongari G, et al. GBA Mutations Influence the Release and Pathological Effects of Small Extracellular Vesicles from Fibroblasts of Patients with Parkinson’s Disease. Int J Mol Sci. 2021;22(4):2215. doi:10.3390/ijms22042215

61. Maloney, et al. LRRK2 Kinase Activity Regulates Parkinson’s Disease-Relevant Lipids at the Lysosome. doi.org/10.1101/2022.12.19.521070

62. Rabia M, Leuzy V, Soulage C, et al. Bis(monoacylglycero)phosphate, a new lipid signature of endosome-derived extracellular vesicles. Biochimie. 2020;178:26–38. doi:10.1016/j.biochi.2020.07.005

63. Wubbolts R, Leckie RS, Veenhuizen PT, et al. Proteomic and biochemical analyses of human B cell-derived exosomes. Potential implications for their function and multivesicular body formation. J Biol Chem. 2003;278(13):10963–10972. doi:10.1074/jbc.M207550200

64. Laulagnier K, Motta C, Hamdi S, et al. Mast cell- and dendritic cell-derived exosomes display a specific lipid composition and an unusual membrane organization. Biochem J. 2004;380(Pt 1):161–171. doi:10.1042/BJ20031594

65. White IJ, Bailey LM, Aghakhani MR, Moss SE, Futter CE. EGF stimulates annexin 1-dependent inward vesiculation in a multivesicular endosome subpopulation. EMBO J. 2006;25(1):1–12. doi:10.1038/sj.emboj.7600759

66. Hsu C, Morohashi Y, Yoshimura S, et al. Regulation of exosome secretion by Rab35 and its GTPase-activating proteins TBC1D10A-C. J Cell Biol. 2010;189(2):223–232. doi:10.1083/jcb.200911018

67. Vieira OV. Rab3a and Rab10 are regulators of lysosome exocytosis and plasma membrane repair. Small GTPases. 2018;9(4):349–351. doi:10.1080/21541248.2016.1235004

68. Bae EJ, Kim DK, Kim C, et al. LRRK2 kinase regulates α-synuclein propagation via RAB35 phosphorylation. Nat Commun. 2018;9(1):3465. doi:10.1038/s41467-018-05958-z

69. Bae EJ, Lee SJ. The LRRK2-RAB axis in regulation of vesicle trafficking and α-synuclein propagation. Biochim Biophys Acta Mol Basis Dis. 2020;1866(3):165632. doi:10.1016/j.bbadis.2019.165632

70. Settembre C, Fraldi A, Medina DL, Ballabio A. Signals from the lysosome: a control centre for cellular clearance and energy metabolism. Nat Rev Mol Cell Biol. 2013;14(5):283–296. doi:10.1038/nrm3565

71. Tsunemi T, Perez-Rosello T, Ishiguro Y, et al. Increased Lysosomal Exocytosis Induced by Lysosomal Ca2+ Channel Agonists Protects Human Dopaminergic Neurons from α-Synuclein Toxicity. J Neurosci. 2019;39(29):5760–5772. doi:10.1523/JNEUROSCI.3085-18.2019

72. Carpenter AE, Jones TR, Lamprecht MR, et al. CellProfiler: image analysis software for identifying and quantifying cell phenotypes. Genome Biol. 2006;7(10):R100. doi:10.1186/gb-2006-7-10-r100

73. Liu N, Tengstrand EA, Chourb L, Hsieh FY. Di-22:6-bis(monoacylglycerol)phosphate: A clinical biomarker of drug-induced phospholipidosis for drug development and safety assessment. Toxicol Appl Pharmacol. 2014;279(3):467–476. doi:10.1016/j.taap.2014.06.014

74. Fernández-Santiago R, Esteve-Codina A, Fernández M, et al. Transcriptome analysis in LRRK2 and idiopathic Parkinson’s disease at different glucose levels. NPJ Parkinsons Dis. 2021;7(1):109. Published 2021 Dec 1. doi:10.1038/s41531-021-00255-x

75. Carpenter AE, Jones TR, Lamprecht MR, et al. CellProfiler: image analysis software for identifying and quantifying cell phenotypes. Genome Biol. 2006;7(10):R100. doi:10.1186/gb-<otherinfo>2006-7-10-r100

76. Dhekne HS, Tonelli F, Yeshaw WM, et al. Genome-wide screen reveals Rab12 GTPase as a critical activator of Parkinson’s disease-linked LRRK2 kinase. Elife. 2023;12:e87098. doi:10.7554/eLife.87098

77. Bebelman MP, Bun P, Huveneers S, van Niel G, Pegtel DM, Verweij FJ. Real-time imaging of multivesicular body-plasma membrane fusion to quantify exosome release from single cells. Nat Protoc. 2020;15(1):102–121. doi:10.1038/s41596-019-0245-4

78. Meneses-Salas E, García-Melero A, Kanerva K, et al. Annexin A6 modulates TBC1D15/Rab7/StARD3 axis to control endosomal cholesterol export in NPC1 cells. Cell Mol Life Sci. 2020;77(14):2839–2857. doi:10.1007/s00018-019-03330-y

