## Supplementary figures and images for "Extracellular vesicle-mediated release of bis(monoacylglycerol)phosphate is regulated by LRRK2 and Glucocerebrosidase activity"

### Supplemental Figure 1

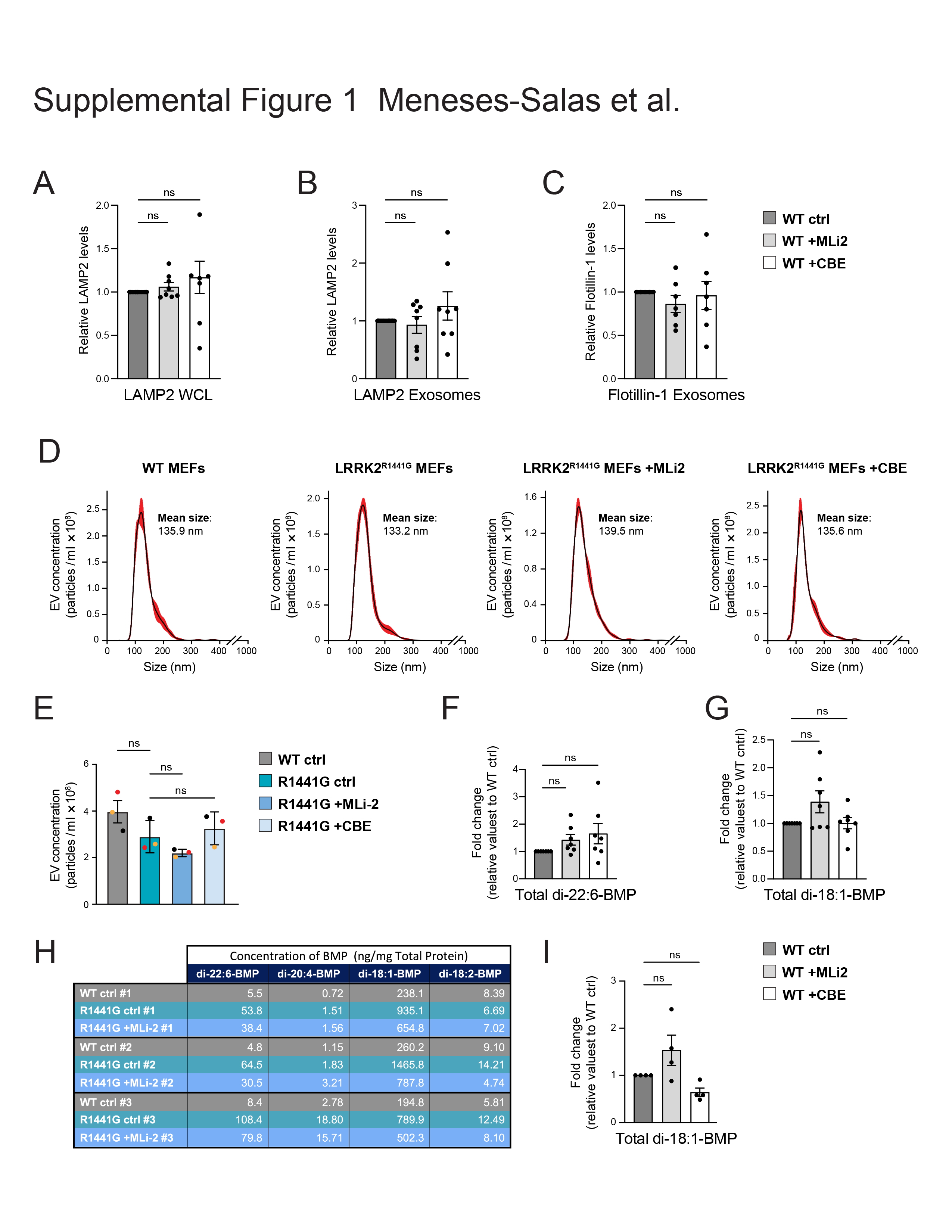

### Supplemental Figure 2

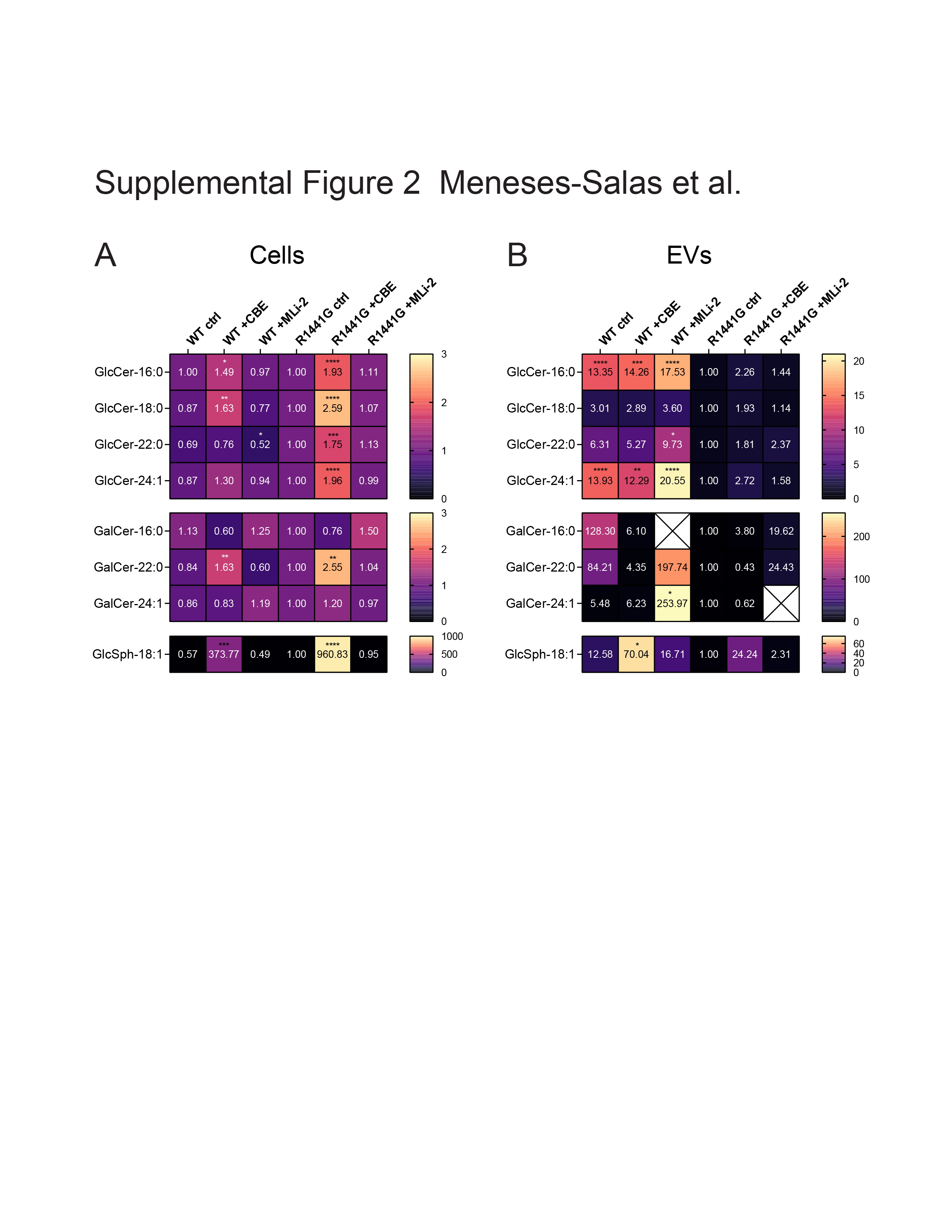

### Supplemental Figure 3

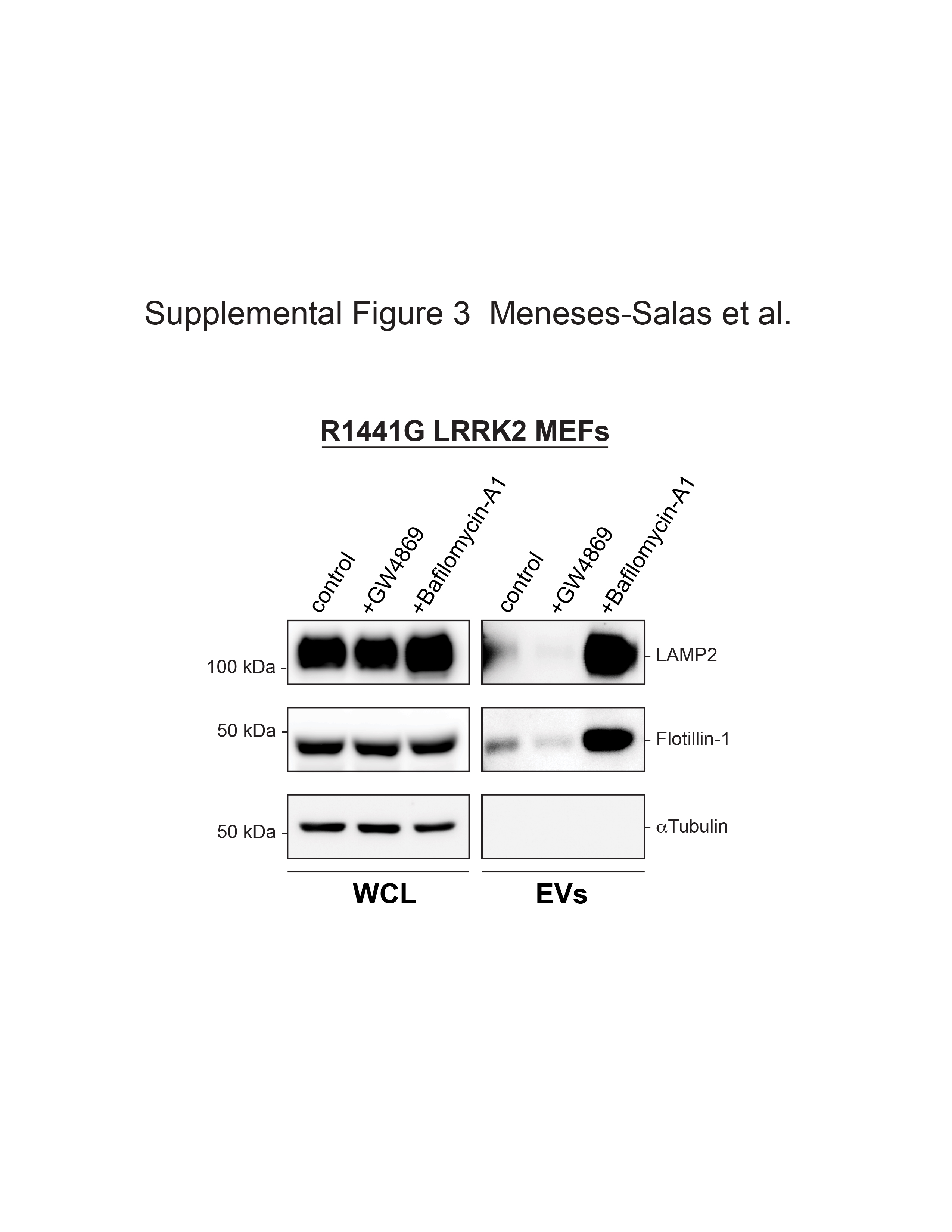

### Supplemental Figure 4

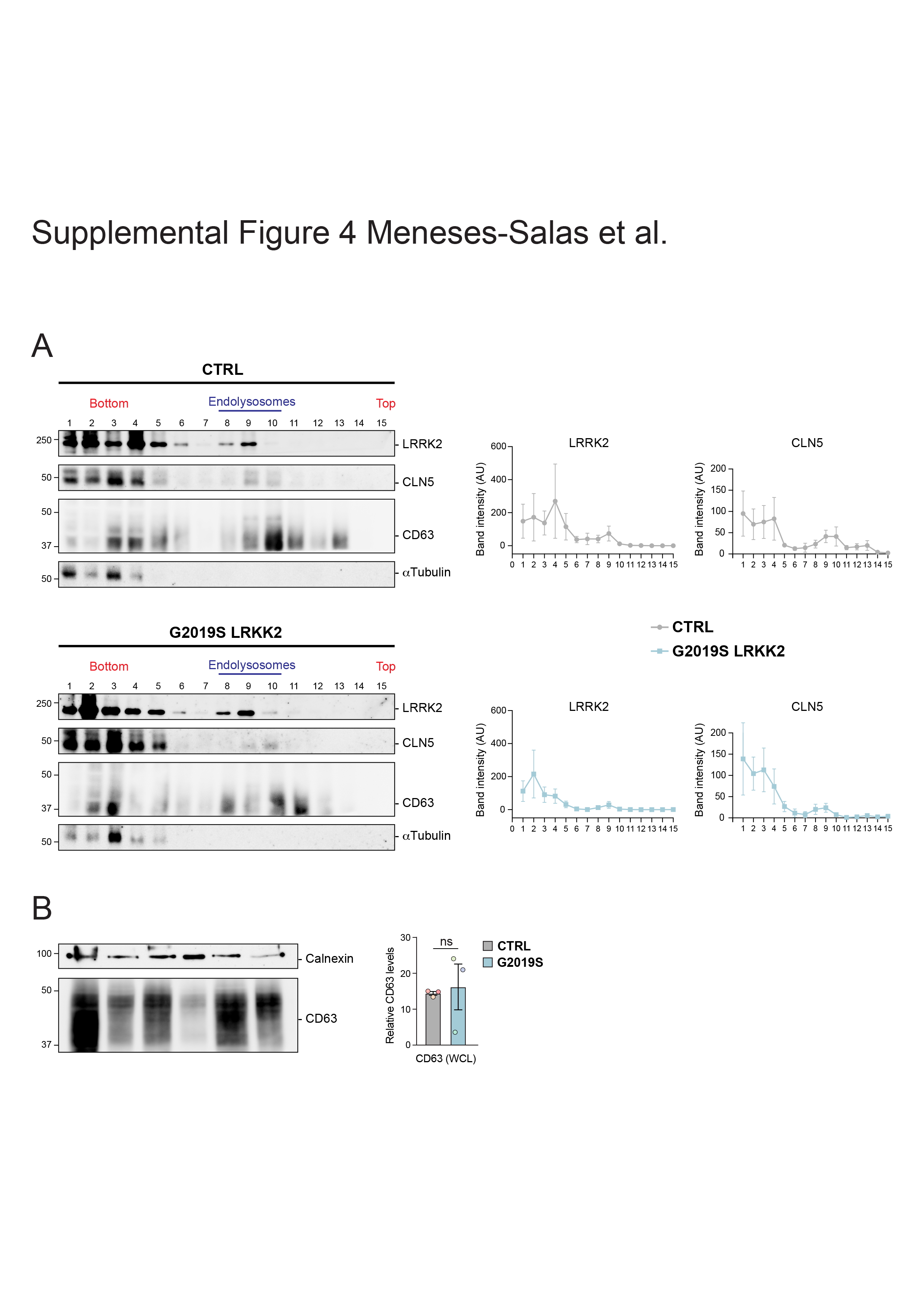
